# “How do we do this at a distance?!” A descriptive study of remote undergraduate research programs during COVID-19

**DOI:** 10.1101/2021.05.17.443632

**Authors:** Olivia A. Erickson, Rebecca B. Cole, Jared M. Isaacs, Silvia Alvarez-Clare, Jonathan Arnold, Allison Augustus-Wallace, Joseph C. Ayoob, Alan Berkowitz, Janet Branchaw, Kevin R. Burgio, Charles H. Cannon, Ruben Michael Ceballos, C. Sarah Cohen, Hilary Coller, Jane Disney, Van A. Doze, Margaret J. Eggers, Stacy Farina, Edwin L. Ferguson, Jeffrey J. Gray, Jean T. Greenberg, Alexander Hoffman, Danielle Jensen-Ryan, Robert M. Kao, Alex C. Keene, Johanna E. Kowalko, Steven A. Lopez, Camille Mathis, Mona Minkara, Courtney J. Murren, Mary Jo Ondrechen, Patricia Ordoñez, Anne Osano, Elizabeth Padilla-Crespo, Soubantika Palchoudhury, Hong Qin, Juan Ramírez-Lugo, Jennifer Reithel, Colin A. Shaw, Amber Smith, Rosemary Smith, Adam P. Summers, Fern Tsien, Erin L. Dolan

**Author notes:** **Author for Correspondence:** Erin L. Dolan, University of Georgia, Biochemistry & Molecular Biology Department, B210B Davison Life Sciences Building, Athens, GA, 30602.

## Abstract

The COVID-19 pandemic shut down undergraduate research programs across the U.S. Twenty-three sites offered remote undergraduate research programs in the life sciences during summer 2020. Given the unprecedented offering of remote research experiences, we carried out a study to describe and evaluate these programs. Using structured templates, we documented how programs were designed and implemented, including who participated. Through focus groups and surveys, we identified programmatic strengths and shortcomings as well as recommendations for improvements from the perspectives of participating students. Strengths included the quality of mentorship, opportunities for learning and professional development, and development of a sense of community. Weaknesses included limited cohort building, challenges with insufficient structure, and issues with technology. Although all programs had one or more activities related to diversity, equity, inclusion, and justice, these topics were largely absent from student reports even though programs coincided with a peak in national consciousness about racial inequities and structural racism. Our results provide evidence for designing remote REUs that are experienced favorably by students. Our results also indicate that remote REUs are sufficiently positive to further investigate their affordances and constraints, including the potential to scale up offerings, with minimal concern about disenfranchising students.

## INTRODUCTION

The global COVID-19 pandemic caused major disruptions to research and teaching across post-secondary education in 2020. Educators and the organizations that support them, ranging from education companies to professional societies to centers for teaching and learning, all scrambled to shift to online experiences for undergraduate programs. A body of knowledge about online instruction, including principles for designing and strategies for teaching online courses synchronously and asynchronously, has been available to inform these changes (e.g., Collison et al., 2000; Means et al., 2014; Palloff & Pratt, 2007). Yet, as STEM undergraduate education has shifted to maximize student involvement in research, a major gap in knowledge has been identified: how to engage undergraduates in research at a distance.

Alternative instructional approaches have been offered up as potential solutions to afford students opportunities to think and work like scientists online or at a distance, such as by analyzing literature or carrying out virtual lab or at-home demonstration laboratory activities (Qiang et al., 2020). Although these approaches are demonstrated to promote student learning and development (e.g., Clark et al., 2009), it is questionable whether they can fully replace the educational value afforded by undergraduate research experiences in STEM. Of particular value is the role that in-person undergraduate research experiences play in facilitating student integration into the scientific community and enabling students to clarify their educational and career interests (Estrada et al., 2011; Gentile et al., 2017; Laursen et al., 2010; Lopatto & Tobias, 2010). Therefore, it was of particular concern that these experiences were relegated to remote experiences in 2020.

Many programs are in place nationwide to offer undergraduate research experiences in the form of internships every summer. One of the longest standing and most widely recognized sources of support for these programs is the National Science Foundation (NSF). This support started in the form of the NSF Undergraduate Research Participation program, which was launched in 1958 (Neckers, 1982). The NSF URP funded projects, known as sites, recruited, selected, and hosted undergraduates as research interns working with faculty mentors and other scientists, including graduate students and postdoctoral associates. Resumed in 1987 as the Research Experiences for Undergraduates (REU) program, REU continues to be one of the largest supporters of undergraduate research experiences in the U.S. (McDevitt et al., 2017). In 2019 alone, 125 sites hosted undergraduate research programs in the biological sciences with NSF support, engaging ∼1,270 undergraduates in research, 68% of whom identified as women and 61% of whom identified as an under-represented minority (Sally O’Connor, NSF REU program officer, personal communication).

About 80% of REU sites funded by the NSF Directorate for Biological Sciences opted to cancel their 2020 summer REU programming due to the COVID-19 pandemic, and 20% – or 25 programs – opted to proceed. In order to document how remote REU programs transitioned to remote research experiences, 23 programs, including one funded by the USDA National Institute for Food and Agriculture, collaborated to generate descriptive accounts of how their programs were designed and implemented. These programs also collaborated with an external evaluation team (OAE, RBC, and ELD) to collect and analyze evaluation data on how undergraduates and their research mentors experienced REU programming, including their perceptions of programmatic strengths and weaknesses and recommendations for improvements. Here we report the descriptive accounts and their alignment with the evaluation results.

Given the unprecedented nature of the situation – specifically, the national shutdown and transition to online instruction by research institutions that host REU programming every summer – we aimed to address two research questions:

- In what ways were summer REU programs implemented remotely or online?
- What were the strengths of these programs as well as suggestions for improvement from the perspectives of undergraduate researchers?

Our results yield preliminary insights into the features of remote REUs that might make them effective for students and their mentors and to inform the improvements of such programs in the future.

## DESIGN AND METHODS

We designed this study to include both observational, descriptive and evaluative components. Through the observational description, we sought to characterize the range of ways REU site programming was implemented during the COVID-19 pandemic. We used a “case series” approach which allowed for the systematic documentation of 23 life science REU programs offered in summer 2020, each serving as a distinct case or implementation of a remote REU site (Grimes & Schulz, 2002). We collected data to document who participated in the 23 remote REU programs, what activities occurred in each program, and when, where, and how each program was implemented. Then, we conducted an evaluation study of the different REU programs from a utilization-focused perspective (Patton, 2008), meaning that we aimed to collect, analyze, and report data that would be useful to REU site principal investigators (PIs). Specifically, we sought feedback from undergraduate researchers and their research mentors on the strengths of the novel, remote experiences as well as suggestions for improving programs both immediately and in future offerings. The results reported here are part of a larger study of remote REUs that was reviewed and determined to be exempt by the University of Georgia Institutional Review Board (STUDY00005841, MOD00008085).

### Programs and Participants

We invited 25 institutions that were involving students in remote or online undergraduate research programs in 2020 to participate in this study. Twenty-three (23) programs chose to participate. The programs were hosted by 24 organizations (e.g., universities, research institutes) in 18 states and 1 U.S. territory and involved 3-39 students and 2-20 mentors per site, with funding from NSF, USDA NIFA, and other sources. One site that was invited to participate in the evaluation did not have the capacity to do research at a distance, so they joined with another site to offer a combined program. Five programs across four sites also involved in-person research experiences for a small number of students, while 21 programs were entirely remote. In this study, we focus primarily on the remote programming and the experiences of students who engaged with their REU and carried out research entirely online. Table 1 provides information about the number and racial, ethnic, gender, and first-generation college status of students who participated in this study.

**Table 1.** Characteristics of students participating in this study. In total, 243 students participated in this study, including 164 women, 71 men, 6 individuals identifying as non-binary, and 2 not reporting a gender. There were 48 students who identified as transfer students and 70 who indicated were first generation college students (i.e., no parent or guardian completed a bachelor’s degree). Students’ racial and ethnic identities are reported, disaggregated by the number of terms (i.e., summer, quarter, or semester) they indicated participating in research prior to summer 2020. Students who identified with multiple races or ethnicities are included in all relevant counts (e.g., a student who reported as Black and Latinx are included in counts for both African American or Black students and Latinx students). Thus, counts may not sum to the totals.

| Race/ethnicity | Prior Research Experience |  |  |  |  |  | Total |
| --- | --- | --- | --- | --- | --- | --- | --- |
|  | None | 1 term | 2 terms | 3 terms | >3 terms | Not reporting |  |
| **African American or Black** | 7 | 7 | 7 | 4 | 11 | - | 36 |
| **Asian** | 6 | 7 | 9 | 8 | 7 | - | 37 |
| **Latinx** | 10 | 14 | 15 | 11 | 12 | - | 62 |
| **Middle Eastern** | - | 2 | 1 | - | 1 | - | 4 |
| **Native American or Native Hawaiian** | 2 | 3 | 2 | - | 1 | - | 8 |
| **White** | 19 | 33 | 35 | 14 | 22 | - | 123 |
| **Not reporting** | - | - | - | - | 1 | 2 | 3 |
| **Total** | 39 | 56 | 61 | 33 | 52 | 2 | 243 |

### Data Collection and Analysis

We collected three types of data. We collected written program descriptions from REU Site PIs, and we conducted focus groups with REU students and their research mentors, as described below. We also collected survey data from students about the quality of their mentorship relationships and the sense of community within their programs, as there was concern that these elements of an REU may be especially difficult to achieve remotely.

#### Written descriptions

We collected written descriptions of each program using a structured template (see Supplemental Materials) to document when, where, and how each program was implemented from the perspective of its PI(s). Specifically, we asked PIs to describe the design and implementation of their program, including program expectations, introductory and culminating events, and weekly activities, shortly after their program was completed. We chose this timing to ensure PIs could describe the implementation of their programs in their entirety (i.e., after all activities were completed) and with accuracy (i.e., soon enough to be able to recollect program activities). We then edited the descriptions to create streamlined, self-similar “program profiles” to allow for quick comprehension and easy comparison of the features of each program. We met briefly with PIs to clarify any ambiguities and fill in any gaps in the profiles before asking for their review, any revision needed, and approval that the profile accurately represented the design and implementation of their programs. Once the profiles were completed and compiled (included in Supplemental Materials), we reviewed the collection to generate a summary description of the REU sites. Site names are included to allow readers to follow up directly with site PIs for details.

#### Focus groups

We conducted focus groups with students in each program at the midpoint and end of their programs. An average of 81% and 67% of students participated in midpoint and end of program focus groups respectively, with percentage by program ranging from 33% to 100% for midpoint and 17% to 100% for end of program. We also conducted focus groups with mentors at the midpoint and end of programs, depending on mentor availability. During all focus groups, we sought feedback about positive aspects of programs as well as suggestions for improvements. For larger programs or instances when not all students were available at the same time, we held multiple focus groups for the program and students chose the one that best suited their schedule. In instances where a student or mentor was unable to participate in a focus group, we solicited responses to the questions by email. All focus groups were recorded to ensure feedback was captured accurately and in its entirety.

The student focus group data were the primary focus of analysis. The evaluation team (OAE, RBC, and ELD) identified strengths for each program and suggestions for improvement by reviewing student responses to the relevant focus group questions. Then the evaluation team created brief, descriptive, and actionable summaries of the strengths and suggestions for each program along with illustrative quotes as supporting data, which were provided in mid- and end-point reports to each program. The evaluation team then carried out an inductive, qualitative content analysis of the reports (Miles et al., 2014; Saldana, 2015). The team independently read each strength and suggestion and ascribed it with a meaning (i.e., to what aspect of the program does this strength or suggestion relate?). The team then met as a group to discuss and refine the meanings, group them into larger themes, and develop definitions of each theme. The evaluation team then carried out a deductive check to ensure that the themes provided a coherent and cohesive representation of the meanings identified across all of the focus groups (Saldana, 2015). Specifically, the team compiled all of the feedback initially identified as fitting a particular theme and reviewed the feedback to determine whether and how it related to the theme. The team revised and refined the themes as needed to ensure they represented a parsimonious interpretation of the data while reflecting the range of feedback identified in the focus groups.

Finally, the evaluation team reviewed all of the reports to identify cross-cutting themes related to the strengths and suggestions and to determine whether each theme was reported as a strength, a suggestion for improvement, or a mixture of the two for each program. In keeping with a descriptive study, our results include detailed descriptions of each program (see Supplemental Material) as well as descriptions of the strengths and suggestions identified through this cross-program analysis. We primarily report on students’ experiences because mentor feedback about strengths and suggestions mirrored feedback from the students.

#### Surveys

To complement the focus group data, we surveyed students at the end of their programs regarding:

- The extent to which they experienced their programs synchronously vs. asynchronously;
- The quality of their relationships with their research mentors (Ragins & Cotton, 1999); and
- The level of connectedness they felt in their programs (Rovai, 2002).

Survey items are included in the Supplemental Materials. Given the research questions and the descriptive nature of the work, survey data were analyzed using descriptive statistics. Means and standard deviations were calculated for the entire dataset as well as at the program level.

Program names have been removed in the reports of the focus group and survey data to protect program confidentiality.

## RESULTS

Here we present the descriptions of remote REU site design and implementation. Then we present the themes that emerged as strengths and areas for improvement during student focus groups. We include survey results to support related focus group findings.

### Remote REU Site Design and Implementation

The REU sites in this study varied in the extent to which the overall design and scientific focus changed to accommodate remote offerings. Some sites shifted to allow students to work in teams with a single mentor or to allow mentors to pair up to work with one or more students. These changes were made for a variety of reasons, outlined in the REU site profiles (see supplemental material). For some sites, restructuring for students to work in teams enabled the involvement of a larger number of students, with groups ranging from two to five students. For other sites, pairing up mentors facilitated the shift to entirely computational projects. For instance, some mentors with bench or field foci of their research paired up with colleagues doing computational work to formulate suitable projects. Some sites that typically had students work in teams dropped the teamwork component of in order to ease logistics. Some sites were already computational in focus and the *Rosetta Commons REU: A Cyberlinked Program in Computational Biomolecular Structure & Design* had been implemented with distributed cohorts in previous years (Alford et al., 2017). For these sites, relatively modest changes were made to accommodate entirely remote participation. Student survey responses indicated that the sites included a mix of synchronous and asynchronous programming (Figure 1).

**Figure 1.**
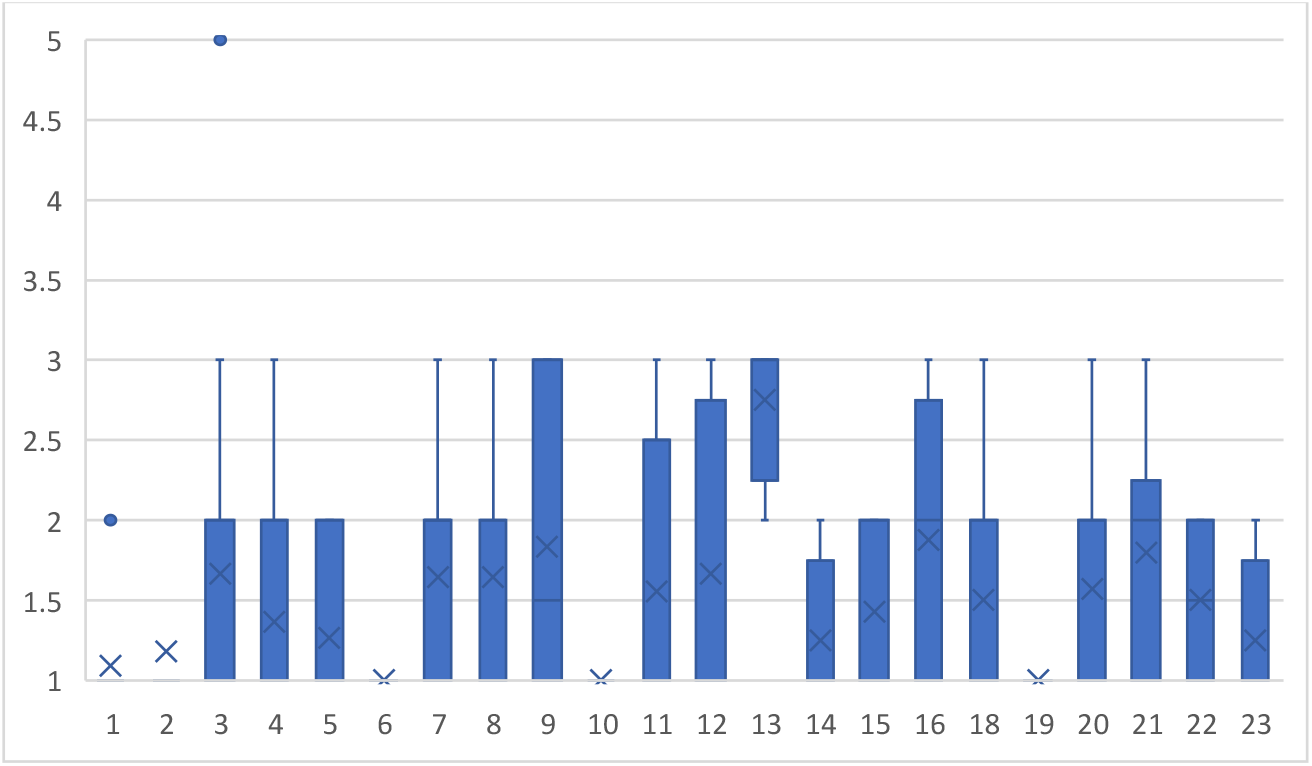
Synchronous vs. asynchronous programming. Students reported that their programs were structured more synchronously than asynchronously (*M=*2.5 SD; *SD=*0.9 with a range of 1= entirely synchronous; 5= entirely asynchronous), with several programs implementing activities entirely synchronously (programs 1, 2, 6, 10, and 19). Lack of consensus in student ratings may indicate variation in how students experienced their programs, with some engaging in more asynchronous activities than others (e.g., watching video recordings of speakers rather than live sessions). Alternatively, students may be perceiving the rating scale differently. Details about the level of synchronous vs. asynchronous programming are provided in supplemental materials. Only data from remote students are included here (i.e., no responses from in-person students in programs 22 and 23).

All sites started with some form of kick-off or orientation for students and/or mentors, although the goals, structure, and content ranged widely. Some sites focused more on social interactions by facilitating get acquainted sessions and community building exercises. Some sites dedicated orientations to building students’ familiarity with the research, the host site, and the expectations for the summer as well as how to go about organizing their remote work. Two sites organized pre-program events or activities, such as discussions with mentors about plans for the summer and how to address issues that might arise as well as workshops to help students get acquainted with research options and begin building computational skills.

In addition to engaging students in research, all sites implemented more didactic knowledge or skill building sessions, either early on or distributed through the summer. These sessions aimed to develop a range of skills, from coding in R to using particular types of software or platforms (e.g., ImageJ, Rosetta Commons, Software Carpentry). Other topics included how to carry out literature searches, navigate databases, use reference managers, apply for fellowships, prepare for the GRE, conduct particular statistical tests, make posters, and communicate scientifically (e.g., writing manuscript-style papers, presenting posters, etc.). All sites included sessions dedicated to the ethical and responsible conduct of research, with some sites addressing particular bioethical considerations such human subjects research and issues related to use of sex and race categories in research (e.g., the *Fungal Genomics and Computational Biology Summer Research* site). The *Exploring 21*^*st*^ *Century Careers in the Biological Sciences: A Comparative Regenerative Biology Approach* site facilitated sessions on innovation, intellectual property, and technology transfer. The *Genes & the Environment REU from Rural & Tribal Colleges* site facilitated sessions on psychosocial skill building, such as managing stress, practicing mindfulness, and engaging in difficult conversations.

All sites hosted panel discussions, scientific seminars, or talks by guest speakers to facilitate students’ professional development beyond research and skill building. Panel discussions addressed a range of topics, from applying to graduate school to offering advice on careers, graduate school, and navigating science as a person of color. Most sites included students in scientific seminars or journal clubs, with some sites expecting students to present their own research in progress or on relevant literature. All sites included at least some discussion about social justice, diversity, equity, inclusion, and/or antiracism. These discussions were facilitated in a variety of ways, from hosting events on antiracism and pride to facilitating movie nights with discussions about the Black Lives Matter and ShutDownSTEM movements.

Some sites included more informal, less structured time in their programming, such as the use of online video communication using Zoom Video Communications, Inc. (Zoom) for lunch hours, coffee breaks, teatimes, and game nights. At some sites, these events were organized by students. Some sites also included Zoom drop-in hours for advice about graduate school, careers, research, technical issues, and troubleshooting. At least two sites collected evaluation data outside of what is described here to make improvements during the summer and identify ways to support students after they completed the program. For instance, the *Bruins-in-Genomics Summer Undergraduate Research Program* site administered regular check-in surveys with students and mentors to identify and address any issues that arose.

All sites ended with a culminating session of some sort, during which students presented their research progress in the form of short talks or posters. Two sites also held award sessions. Talk formats ranged widely from 10 to 15-minute individual or team presentations followed by a few minutes of questions and answers, to 3-minute thesis style presentations or other very short talks. All sites required students to produce one or more products, such as posters, talks, papers, proposals, or videos. The *Cary Institute of Ecosystem Studies REU* site required students to generate “data nuggets” (http://datanuggets.org/), which are mini-research projects or tasks that can be used in K-16 instruction to develop students’ science research skills. Some programs made a point of encouraging students to invite family and friends. The *REU Site at The Morton Arboretum: Integrative Tree Science in the Anthropocene* included keynote speakers of color. The *Rosetta Commons REU* site held their culminating event as part of a larger conference being held by the Rosetta Commons community (https://www.rosettacommons.org/). The *Training and Experimentation in Computational Biology* site held their closing poster session in virtual reality.

### Strengths and Areas for Improvement of Remote REU Sites

Students in this study described the strengths and areas for improvement of their remote REU site in terms of 10 overarching themes (Figure 1). Three themes that emerged as strengths across sites were the (1) quality of mentorship, (2) opportunities for learning, and (3) sense of community within labs and programs. Two themes that emerged as primary areas for improvement were the (4) cohort experience and (5) unstructured nature of research and remote work. Two themes emerged as having both beneficial and problematic elements: (6) program logistics and (7) opportunities for professional socialization. Finally, three themes were identified less frequently across programs and were experienced as either strengths or areas for improvement depending on the site: (8) networking, (9) technical issues, and (10) diversity, equity, inclusion, and justice. Each of these themes is defined and described below in numerical order.

#### Theme 1. Mentorship: Students described the quality of support they received from their research mentors to help them learn, make progress in their research, and be successful in their programs

The main strength across most of the sites in this study was the quality of the mentorship. Students in 15 sites emphasized the quality of the mentorship they received, in terms of technical and career support as well as psychosocial support. One student described in detail the mentorship they received:

The mentor that I had personally, they went out of their way to make sure I was in a good area or ask how I was doing. My mentor in particular was [having a personal situation]. So he had to leave for a while. I had a technician of his take over and she was amazing as well. Even while his family was going through that he would message me to see, ‘How are you doing? How’s your research going? Is there anything that I can do?’ It was going above and beyond to make sure that I was understanding what I was doing and getting the most out of this experience.

This quote captures a sentiment expressed by other students – that mentors provided both direct support and indirect support by connecting them with someone who could help when the mentor was unable to do so. The mostly positive experience students had with their research mentors is also evident in their overall positive ratings of the quality of their relationships with their mentors (Figure 2).

**Figure 2.**
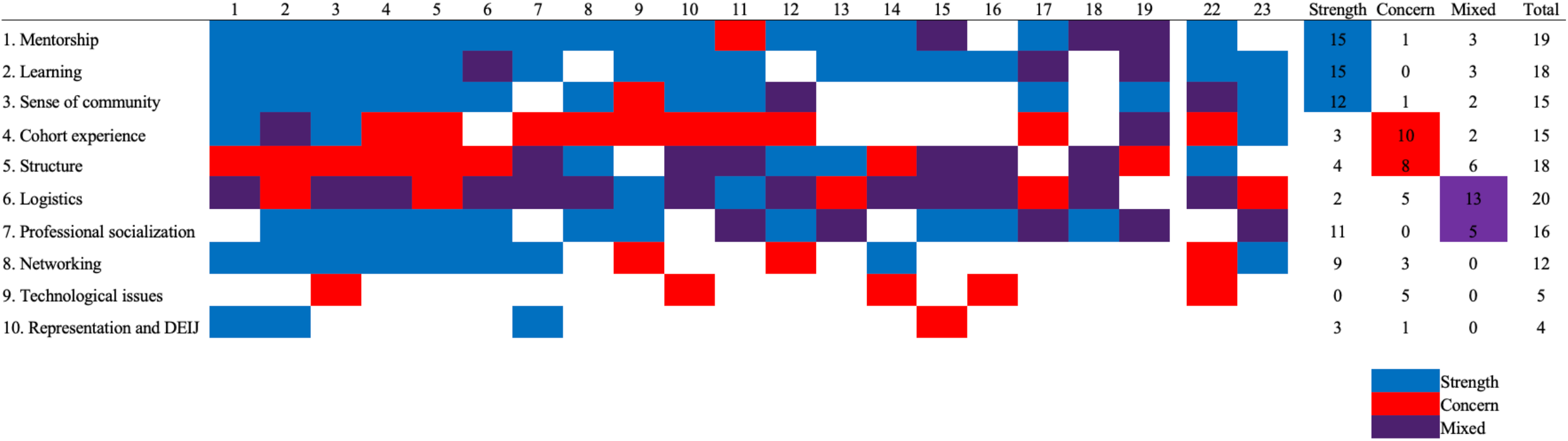
Student-identified strengths and areas for improvement in remote REU sites. This figure provides an overview of the strengths and areas for improvement for each of the 21 programs in this study, which are numbered across the top. Programs 20 and 21 are not included here because students in these programs did not participate in focus groups. Programs 22 and 23 are separated because they included substantive in-person components. Blue indicates that the areas of strength (three most common in the top three rows). Red indicates areas in need of improvement (next two rows). Purple indicates a mixture within a program with some students emphasizing this as a strength and others as an area in need of improvement (next two rows). The bottom three rows feature themes that were mentioned by students in fewer programs. The four columns on the right are sums of how many programs had students reporting the theme as a strength, a concern, or a mix, with the total indicated how many programs had students commenting on the theme regardless of whether it was a strength or concern.

Students across sites noted how their mentors forged connections between them and the rest of the research group so they could reach out and ask questions. One student noted that “it is helpful knowing if I get stuck on something, (my mentor) is available.” Several students noted that they appreciated their mentor’s ability to strike a balance between providing support and allowing students to answer their own questions. One student noted that their mentor “[made] sure [they were] on track. It wasn’t too overbearing, but they were also always making sure I was going along on the project.” Another student described how their faculty mentor was open to feedback such that, when the student expressed concerns about how their experience was going, “it actually improved once I talked to my PI about what was going on and what I needed from her, which helped. That made a big difference.”

Students also noted the ways that mentors provided psychosocial support. Most students who commented on mentorship felt that their mentors cared about them not just as a scientist, but as a person. For instance, one student appreciated that their mentor “was really invested in them and invested in their research.” Another student noted that their personal relationship with their mentor is “something [they] cherish a lot.” Students also appreciated how mentors were responsive to how the pandemic could be affecting their work. One student observed “there are so many assumptions that can be made about the students,” and students appreciated mentors’ willingness to be flexible around complications that arose from working from home. Finally, students repeatedly mentioned how mentors quelled their anxieties around asking for help and that their mentor “never make [them] feel dumb for needing help.”

Students in one site indicated that the mentorship they received was inadequate and students in three sites had mixed experiences with mentorship (see outliers in Figure 2). In these instances, students expressed concern that the time they were able to spend with mentors and the ways they were able to communicate (or not) with their mentors compromised the quality of their experience. For instance, some students who were struggling to make progress on their project felt they could not just “drop in” to ask a question or get help. They perceived that their mentors would have been receptive to providing drop-in help if the program had been in person, but they didn’t see a way to accomplish this remotely. One student indicated they had a set weekly meeting with their mentor and otherwise weren’t “allowed” to contact their mentor with questions except for emergency situations. This often meant that they would reach an impasse in their research and be unable to make progress until the next weekly meeting.

One point was made during a mentor focus group that was not otherwise represented in the student results. These mentors explained that the remote nature of the REU program made it more difficult to oversee and manage what students were doing on an hour-by-hour or day-by-day basis however, they were pleasantly surprised by how much students could achieve without oversight. In other words, the circumstances made it such that mentors were by default more hands off, which resulted in students having more autonomy to make decisions and solve problems on their own. The mentors in this group planned to apply what they learned to their in-person mentoring relationships by giving students more freedom to make progress and decisions on their own.

#### Theme 2. Learning: Students described gains in knowledge, skills, or abilities as a result of participating in remote research

Students in 15 sites emphasized how much they learned from their research experience. Students reported gaining knowledge in the content area of their research and vastly improving their coding skills; one student describing their coding abilities as “phenomenally improved.” Even for sites where computational biology was not a major emphasis, the remote nature of the research meant that students carried out projects that involved coding to query datasets and conduct analyses. Students perceived that their research experiences provided a “real-life” context for learning to code, which was superior to learning coding through coursework, as one student noted: “be[ing] able to actually use it in a project was so much better for learning how to program than anything I could have learned in a class at my university.” In addition, students perceived that their new skills would be “so beneficial for future research and future labs.”

Beyond gaining content knowledge and technical skills, students reported learning more about the research process and gaining confidence in their own abilities to be successful in research. One student noted that “when [they] first started,” [they] thought it would be super hard to conduct research, and it was difficult, but it’s not as unattainable as [they] once thought it was.” Beyond this, students report developing other professional and scientific skills such as troubleshooting. Several students gained a new appreciation for solving problems on their own, expressing that “figuring out things for yourself has become satisfying” and that they now felt “equipped with the skills to be able to troubleshoot problems when I have them.” Students expressed surprise that they were able to grow in their knowledge, skills, and confidence in such a short time while working remotely, one student explaining that “[at first, I was] really nervous putting things together… but toward the end I was really communicating with my colleagues.”

#### Theme 3. Sense of Community: Students described the sense of being connected to and comfortable with their research groups, sites, or broader scientific communities

(Note: Students described their sense of community as distinct from being part of an undergraduate research cohort. Thus, cohort experience is described separately below.)

Students in 12 sites emphasized how their sites and their research groups created a sense of community, which manifested in a variety of ways. For example, some students described how their sites created a culture where students felt they could “go to anyone for help” and that this environment allowed them to “see how collaborative research really is.” Some sites and research groups ensured that students had ample opportunities to interact with graduate students other than those who served as their research mentors, and that this had a “profound impact on [their] overall experience” and “play[ed] a big role in feeling welcome to [their] lab group.” Students emphasized the importance of making these connections early in the summer so that it was easier to seek out that guidance later in the program. Yet another student noted that the level of engagement by everyone involved in the program helped them feel like part of a community. The student described that, during presentations, “everyone is really supportive and engaged and they give you really valuable feedback, not just for the sake of giving feedback, but because they’re actually engaged with what you’re saying.”

The sense of community students developed is also evident in their overall positive ratings of their connectedness with their sites (Figure 3), although students were less favorable about this than about their relationships with their research mentors. Students in one site expressed frustration that there wasn’t transparency about whether they could seek help from others outside their research group or what resources were available to provide help. They explained that there was a “resource sitting there for everybody and only a select few knew about it.” In this instance, it appeared that one or a few research groups made their students aware of the resource but that other research groups and the site administrators did not, which created inequity that undermined the sense of community in the program. Similarly, in this program, certain research groups made an effort to connect their students with other faculty. These students appreciated the opportunity to develop relationships with faculty members other than their mentors and to become part of a “community of different scientists.” Students who did not have this experience were eager for it, indicating they wanted to learn from a broader and more diverse group of faculty members about topics beyond “research and what they look for in graduate students,” such as “how they became a scientist and what they see as lab culture.”

**Figure 3.**
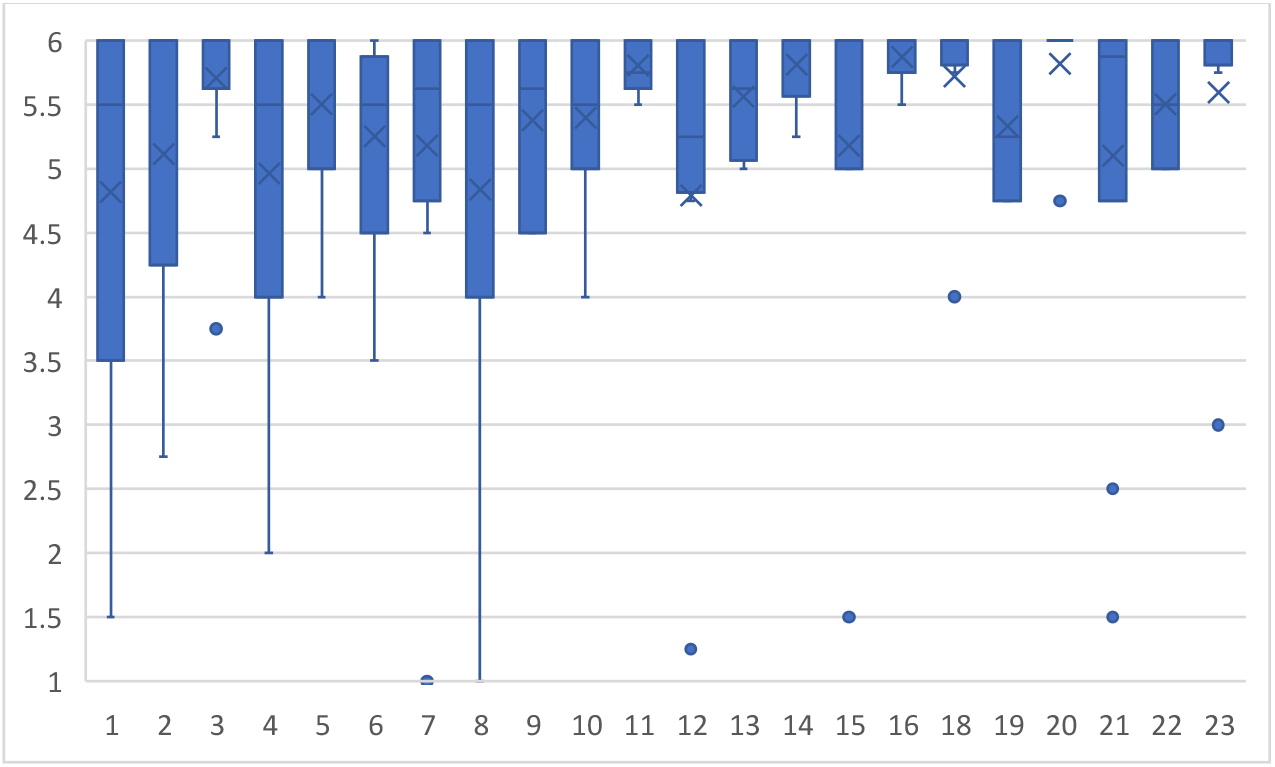
Relationship quality. For the most part, students rated their relationships with their mentors quite positively (*M*=5.3 out of 6; *SD*=1.2). This figure shows student ratings by site, with 6 indicating strong agreement and 1 indicating strong disagreement with a positive statement about relationship quality (see supplemental materials for items and rating scale). The X signifies the site mean and the bar indicates the site median. Some negative ratings were observed, which reflects the mixed or negative experiences of a small number of students. Only data from remote students are included here (i.e., no responses from in-person students in programs 22 and 23).

#### Theme 4. Cohort experience: Students described the sense of being banded together as a group of research interns, feeling close to and engaged with other undergraduate researchers in their cohort or feeling isolated or disconnected from the group

Students in 12 sites indicated that they missed feeling like a cohort and expressed concern about missing out on a cohort experience. In one of these sites, students reported mixed perceptions of cohort feelings, with some finding it easier and some finding it more difficult to get to know one another. One student expressed this mixed feeling in describing their end-of-program poster session, noting that “it was sweet to see the other interns and to like want to go to their [Zoom breakout] rooms and just check in on everyone. I still feel like, even though [the program] wasn’t in-person, it built camaraderie and a cohort.” Across the 12 sites, students reported several factors that prevented or undermined the development of a cohort feeling. First, some sites involved only a few students. Students thought that the small number was insufficient to provide a cohort experience. Second, at least one site held fewer whole group events as the summer progressed to allow students to focus their attention on their research. Students in this site indicated that they would have preferred to continue meeting weekly as a whole group to continue to get to know each other. Finally, students found it difficult to have more casual interactions that normally occurred when working alongside others. They felt that this limited their abilities to network and build relationships with other students. One student explained that their site “tried to do little things to build community for the students who were remote learning, but it as far as I can tell, it kind of fell short, I was really only communicating with the people in my [research] team.”

Other students lamented the loss of informal interactions because they were not “able to ask a neighbor ‘hey, can you help me out with this?’” One student explained how not getting to know people on a personal level meant that they were not able to alleviate some of the nervous feelings associated with asking questions. Students had mixed feelings about social hours on Zoom for cohort building. Some appreciated having game nights or other social activities (e.g., Pictionary on virtual whiteboards, bingo, escape room, trivia night, Jackbox, virtual meditation or yoga), while others felt “Zoom fatigue” after many hours of program and research activities on Zoom. Students in several sites suggested integrating cohort building into regular workweek activities rather than as an additional activity. For instance, students in several sites expressed the desire for synchronous, online work time on Zoom to simulate an in-person collaborative work environment. Students could join the call and ask impromptu questions or talk through ideas as they worked. Similarly, students wanted to use GroupMe or Slack among themselves to communicate about non-research related things and get to know each other.

Students in three sites indicated that their sites supported a sense of being part of a cohort of undergraduate researchers. These students emphasized that they still felt a sense of connection with other undergraduate researchers in their site despite the remote circumstances. They appreciated the opportunity to interact with other undergraduates and they reported that doing activities as a group and being encouraged by site leadership to socialize among themselves helped to achieve this. Other factors that promoted their sense of camaraderie in their cohorts included talking about things “outside the scope of our respective projects,” such as students’ roles in the broader scientific community and in the world given the country’s raised awareness of systemic racism and racial injustice. For instance, one group of students commended their site for making time and creating a safe space for discussion about BlackLivesMatter and ongoing racial injustice in honor of the #ShutdownSTEM initiative. This group reported that these activities have helped to both “build a dialogue about the issues and build a community” among the cohort. Students in another site appreciated the intentionality displayed by their site’s leadership to facilitate a sense of community. This site established a committee structure, which gave every student a way to be involved and promoted a sense of inclusion. Several students indicated that having a social committee helped to enhance the cohort experience. Students also noted that having a student-only GroupMe group or Slack channel as well as the use of smaller breakout groups on Zoom all facilitated getting to know one another and promoted a cohort feeling.

#### Theme 5. Structure: Students described program design elements, such as schedules, workflows, expectations, milestones, or deadlines, which helped them organize work and manage time

Students in 14 sites indicated that they were struggling with the lack of structure inherent to remote work and to research. They noted that having scheduling flexibility was helpful because their circumstances were so unpredictable, but that the extent of the flexibility was “daunting” and made time management difficult. They expressed concern that they didn’t know how much progress they were expected to make each day, and they struggled to define when their workday should start and end. The lack of clarity regarding how much to work and what was expected of them left some feeling like they had “to work on their project at all times” and prompted some to work longer hours. For others, they felt as though they had extra time that could have been used more productively. If they had been onsite, they would have sought additional things to do, but they weren’t sure how to do this at a distance. Having mentors with more of a “hands-off” approach exacerbated these issues.

Students across sites made several suggestions for adding structure that would have allowed them to better gauge whether they were on track in their research and programs, including:

- Defining a daily or weekly schedule or offering suggested schedules, including expected number of hours per day (even “clocking in”) and whether and how much they should take breaks to prevent burnout,
- Defining “checkpoints,” “check-ins,” “assignments,” or “intermediate goals” throughout the program to help with gauging progress and avoid tasks “hitting [them] all at once” at the end,
- Ensuring mentors set aside time every day or two or schedule standing meetings to provide guidance and instruction,
- Requiring students to write brief weekly updates or reports for their mentors to check to ensure they are making sufficient progress,
- Scheduling midpoint progress meetings to get feedback from mentors about the progress they have made, the quality of the work they have completed, and goals and potential improvements for the remainder of the summer,
- Providing a list of optional tasks or recommendations for what students could be doing if they had extra time, such as additional reading, writing, or analysis tasks, working on other projects when they have downtime on their main project, and additional skill building, and
- Hosting one or two sessions with mentors or site leadership to share how they manage their workdays and brainstorm strategies for time management (e.g., what to do, in what order, and when to get things done by) and structure that helps them to “organize their day, set priorities, and meet goals.”

Some of the students who made these suggestions thought that increased structure would not only help them better gauge their progress, but would also help them avoid distractions and “set firmer boundaries with family members during times they have set aside for working.” Some students shifted to creating their own structure to mitigate the lack of structure inherent to working from home, including “making a daily checklist…that motivated me to get things done in the day” and “mak[ing] a [physical] workplace that’s separate from where you rest, just so you can separate working life better.”

Students in four sites indicated that their sites provided important structure to help them stay on track throughout the summer. One site required students to prepare a research proposal and complete other mandatory assignments, which helped them “refocus” and “make sure (they) knew what (they) were talking about.” They explained that “the more mandatory assignments [they] had, the more on track [they were] because they had to force [themselves] to reevaluate [their] understanding and application [of their knowledge and skills].” Other sites had regular meetings with site leadership, such as start-of-week check-ins, that ensured they set goals and gauged progress on a regular basis and got feedback and help before too much time had passed if they were off track.

#### Theme 6. Site logistics: Students described operational aspects of sites, including onboarding, meetings, communication, and pacing, which improved or undermine their experience

Students in 15 sites indicated that several aspects of how their sites operated made it possible to navigate the program smoothly at a distance. These aspects included frequent meetings with their mentors, their cohort, and/or the site leadership, clear and open communication between students, mentors, and site leadership, and proper program pacing. Students reported that the inclusion of frequent meetings, such as daily with their mentors and weekly meetings in their sites, helped them to stay focused and motivated and to feel connected with others in the community despite being physically distant from them. They also noted that these meetings made communication easy to maintain and were important for their success in the site, helping them “feel a little bit more connected and less on my own.” Students also noted that regular communication in advance, such as weekly announcements of upcoming events and other key information, made it easier to ensure they were in the right places at the right times and had sufficient time to plan their research around site activities. Students appreciated having access to this information in a single location or platform so they could find it when they needed it. Students in several sites commented that their sites started more slowly, helping them acclimate to working online at a distance and get up to speed on their research. They also appreciated that pacing changed over time, allowing more time as the summer progressed to focus more on research and less on site activities.

Students in 17 sites commented that some logistical elements were missing, which compromised their overall experience. Examples included poor or sporadic communication, uneven program pacing, and difficulties with onboarding. Regarding communication, students reported wanting more open and consistent communication among them, their mentors, and site leadership. For instance, some students reported getting announcements on multiple platforms, which led to confusion about where and when to find needed information. In some instances, announcements came with such short notice that students missed activities. Other students expressed concern that their mentors seemed unaware of site activities, which resulted in site activities feeling separated from or in conflict with their research activities. In these instances, students felt like they had to choose between their site responsibilities and furthering their research. Students suggested that summer program calendars be shared with mentors in order to alleviate confusion. They also suggested scheduling events at a particular time and communicating these times with mentors and students sufficiently far in advance to allow for planning. Students indicated that mentors needed to seek mentee input when scheduling meetings since everyone had different schedules, often in different time zones.

Students in multiple sites struggled with the pacing of their program. They expressed concerns about pacing both within a day and across the summer. Day-to-day, students emphasized the importance of limiting the number of online meetings and sticking to schedules rather than letting meetings run over time. Across the summer, students indicated that site activities should be more evenly spread throughout the summer, rather than front-loaded at the beginning. This change would allow for more time to start research and enable just-in-time guidance and support, such as writing workshops when students would be writing instead of early in the summer. Finally, given the remote nature of the sites, students needed functional computers, software, and network access as well as institutional credentials to access institutional resources and functions.

#### Theme 7. Professional socialization: Students described how sites helped them gain insight into graduate education and research careers and to envision themselves pursuing further education and careers in science

Students in 15 sites indicated that their sites facilitated their professional socialization despite the remote circumstances. One approach that sites used to accomplish this was to host online sessions related to graduate education, including webinars about fellowships and funding opportunities, panels with current graduate students, and workshops for GRE preparation^1^. Students found it inspiring to hear from current doctoral students and the many different paths they could take to graduate school. One student highlighted how an NSF grant workshop was so “motivating” that it “inspired [them] to get [their] academics in order [so that they could] get research opportunities in the future, and eventually get to graduate school.” Several students noted that these sessions served as a “mental health break” from the challenging work they were doing in their research.

In addition to engaging students in research, sites supported students’ professional socialization by hosting sessions highlighting the diversity of research careers. Typically, these sessions involved panels of scientists from a wide range of fields, careers, and backgrounds, providing students insights into “what it’s really like to be a researcher, the good and the bad,” and helping them to discern whether they would like to pursue a career in research. Students noted that a major advantage of online panels was that they met scientists from a wide variety of fields from all over the country, which they thought might not have happened if the site was in-person. Some students felt their sites could have done more to integrate them into the research community. Typically, these sites did not offer workshops related to graduate school preparation or had limited if any interactions with speakers, panelists, and other students.

Through attending workshops about graduate school, hearing from current doctoral students and scientists during panels, and doing research, students reported feeling that they had “found their purpose.” For instance, one student indicated that “I live close to [a Native American] reservation, and I’m a [member of this tribe], too. It was hard to not be able to do anything for my people [during the pandemic]… I didn’t know how to help out. When I heard about this research experience, it was like, ‘Hey, this is how I can actually help in some way.’” More generally, students also commented developing “confidence in [themselves]… and what kind of research [they] want to do” and “reassurance that [they] can do this and that this is something that [they] can see [themselves] pursuing.”

#### Theme 8. Networking: Students described opportunities to meet and build relationships with others who may be helpful for learning and career development

Students in six sites explained how their sites provided opportunities to meet and build relationships with faculty, professionals, graduate students, and peers who could help them learn or otherwise advance toward achieving their education or career goals. Several students felt that they had plenty of opportunities to “expand their network.” For some, networking mitigated the feeling of being isolated, explaining, “if we didn’t get to meet as many people from [the institution] as we did, the [remote] experience would have been significantly more isolating.” In fact, some students commented that “the most impactful” thing they got out of their research experience were the connections they made throughout the summer, as one student describes, “The community was something that was really helpful for me, especially looking at the network of resources and the networks of labs to join for possible next steps in my future as well as the future of my research.” Several students expressed how grateful they were to finish their program feeling like they had met people who could help them as they progress in their careers. One student commented that, before their experience, they didn’t realize how collaborative the scientific community was and thought that it was “really awesome to see that, from this one opportunity, [they] now have connections to [so many] different places.”

Even in programs where students noted networking was a strength, this varied by lab group, with some groups fostering more connections than others. For example, several students commented that they heard from their peers about interacting with graduate students and they wished they had more opportunities to do so. Students also expressed a desire to develop relationships with faculty other than their own mentor. They felt they had learned so much from their own mentor, that their experience could only be enhanced by learning from other mentors. Some specifically wanted to hear from faculty members about topics “beyond research,” such as “how they became a scientist and [how they view] lab culture,” and these students mentioned that having meet-and-greet hours with faculty would be an impactful way to facilitate these connections. Other students suggested having their work reviewed by more than one mentor would afford opportunities to get more feedback and build rapport with other mentors. Students acknowledged that they felt personal “responsibility to network and make those connections” as well as a responsibility of the site to facilitate building relationships, especially given how challenging this was for students to do remotely.

Students indicated that sites supported networking in multiple ways. Some sites encouraged students to talk and work with lab groups and mentors other than their own. Other sites took advantage of the remote circumstances to organize cross-site activities and events. Students who participated in these opportunities appreciated connecting to researchers both within and beyond their site and were grateful that this enabled them to be able to work with mentors with expertise in their research interests. Students in some programs had the opportunity to help choose speakers and organize seminars. One student explained that this was an advantage of a remote site because they had “a wider range of speakers because we can reach people all over the world right now,” and how “hearing from a researcher in [another country] was especially exciting.” Having informal settings for interaction was another tactic that supported networking. For instance, one site had weekly check-ins with the directors, which one student indicated was a favorite part of their program.

#### Theme 9. Technological Issues: Students described issues with technology that undermined or limited their experience

Students in five sites reported several issues with technology that compromised their research progress and their overall experience. First, some students had difficulty accessing communication platforms (e.g., an institutional learning management system) either because they did not have the appropriate credentials for access or because the platform itself was “confusing to navigate” or “hard to use.” Second, some students described how their sites used multiple communication platforms, which made “easy to miss things” when certain events or activities were announced in one platform, but other key information was available in a different platform. Third, some students did not have sufficient internet connections or access to a computer with sufficient computing capacity or credentialing to allow for access to necessary software. These issues were identified by sites and PIs were responsive to student needs, yet the time it took for technology issues to be solved ultimately limited the amount of progress students felt they could make in their research. Finally, some students indicated that they did not have enough support with coding or learning to code. Several of these students explained that, by the second half of their programs, they had found someone that they could ask for coding help when needed. Yet, they wished these connections had been made available to everyone in the program early in the summer so that they had equal access to support and could have made better progress throughout the summer.

Interestingly, no students indicated technology as an area of strength for their site, possibly because students expected technology to work and thus only noticed when their expectations were not met. Students who reported having technology issues made three suggestions for preventing these issues or mitigating their impacts in the future. First, they recommended selecting a common, easy-to-use platform for communication such as group messaging (e.g., GroupMe, Slack) or email lists. Second, they recommended setting up institutional credentials and conducting technology audits in advance of the site start date by determining the technological needs of each research project and the computing and internet capacity to which each student has access. If the needs exceed the capacity, there should be sufficient time to ship suitable computers (this was done by the *Summer Integrative Neuroscience Experience in Jupiter* at Florida Atlantic University), set up improved internet access, and ensure students have needed credentials in place. Finally, they recommended making transparent to all students the individuals who could provide coding support. This support could be provided by the research group, the site, and/or the institution, depending on needs and resources.

#### Theme 10. Diversity, equity, inclusion, justice, and representation: Students described how sites created time and space to discuss social justice topics

A review of the REU site profiles (see Supplemental Materials) shows that all sites facilitated at least one formal or informal discussion or event regarding diversity, inclusivity, social justice, or anti-racism. However, students in only three sites mentioned this aspect as a strength of their site. One possible explanation for this is that many of these events and discussions were informal in nature or limited in scope so students might have not perceived these discussions as a formal part of the site or sufficiently substantive to be mentioned during the focus groups.

Students in two sites spoke about how their sites set time in their schedules to discuss issues around diversity, equity, inclusion, and social justice, as well as representation of individuals from backgrounds that are traditionally excluded or marginalized from the sciences. Students in these sites noted that the discussion of the larger national social justice conversation made them feel as though they were “people and not just scientists.” These students also appreciated the opportunity to bring their whole selves to their research experience and they appreciated being encouraged to “talk how they like to talk.” One student explained that offering remote REU experiences allowed for participation in research by people with disabilities or other circumstances that prevented traveling to a distant site. One student indicated that they had not previously imagined applying to graduate school but found it “inspiring” to hear from graduate students who took non-traditional paths to graduate school.

In one site that held multiple events related to diversity and inclusion in STEM, students explicitly highlighted representation and DEIJ as an area of weakness due to the absence of people of color in workshops and seminars. Additionally, they mentioned that they would have appreciated receiving advice from individuals from more economically diverse backgrounds and diverse career paths “other than ‘went to undergrad, went to grad school, got a job, paid off my loans.’”

## DISCUSSION

When considered collectively, these results indicate that remotely implemented REU sites can, at least under certain circumstances, afford many of the same opportunities that in-person sites offer. Students indicated that they learned, experienced quality mentorship, grew professionally, and expanded their networks. They felt like they became a part of a research community that would not have been available to them if they had not participated in remote research. This finding adds to a previous report that students in a mostly remote REU site were able to develop a sense of community (Alford et al., 2017). In addition, the remote implementation of research experiences appeared to provide access to networks that might not have otherwise been available. Specifically, the remote implementation prompted sites to invite individuals from all around the country and even around the world to meet with students as speakers, panelists, and collaborators, thereby expanding students’ connections far beyond what might have occurred in-person. These results should provide some reassurance that remote REUs are worth offering and may offer some advantages over or in addition to in-person programming. For example, remote sites could involve undergraduates in research whose personal situations would preclude participating in an onsite program. In-person sites could consider adopting some of the strategies used during remote programming, such as networking across sites and holding sessions using video conferencing so that students can interact with speakers, panelists, and collaborators beyond those who are available onsite.

Our results also indicate that several elements of REUs were more challenging to implement at a distance. For instance, even though most students reported experiencing quality mentorship, others indicated that their mentorship experiences fell short of meeting their needs. In these instances, students perceived that the absence of quality mentorship stymied their research progress and professional growth. It may be that the quality of mentorship simply varies within sites, which is consistent with research on mentorship in undergraduate research (Byars-Winston & Dahlberg, 2019; Hernandez et al., 2017, 2020; Limeri et al., 2019). Alternatively, some mentors may be less prepared to provide support at a distance and may need additional guidance and support on how to do so effectively. There is little if any research on how to prepare mentors to remotely support undergraduate researchers, which presents the unique challenge of not being able to “drop in” to see how an undergraduate researcher is doing or otherwise engage in informal interactions that are critical components of effective mentorship (Ragins & Cotton, 1999). However, sites can put several measures in place to avoid or mitigate the impact of insufficient or problematic mentorship, which are consistent with recommendations from the National Academies on effective and inclusive research mentorship (Byars-Winston & Dahlberg, 2019). First, sites can set clear expectations for the frequency with which mentors should be expected to communicate with students and the flexibility of that communication. Second, sites can collect data on mentorship support and quality and determine whether certain individuals are not well suited to mentor students at a distance or in general. Finally, sites can conduct midpoint checks with students about the mentorship they are receiving, including what is working well and what needs to be improved. This feedback can then be used to help mentors and students improve their mentoring relationship or remove students from situations that are deemed sufficiently problematic.

Although students reported developing a sense of community with their research groups, they expressed concern about missing out on being part of a cohort. This concern was mitigated somewhat by sites that promoted informal interactions and at least one site that made use of a committee structure through which social activities were promoted and each student had a specific responsibility as part of the site. This is consistent with research on community building, which indicates that community can be fostered through shared tasks (Kim, 2006; Lave & Wenger, 1991; Wenger, 1999). Students in remote sites shared research tasks and thus built community with their research groups. For the most part, however, they did not have shared programmatic tasks. Although it is not clear that in-person REUs have shared programmatic tasks, it may be that ad hoc, informal interactions that occur in in-person sites promote identification with the group and shared responsibility for its growth and success. The site that made use of a committee structure was able to promote cohort building even at a distance. Other sites could consider establishing roles or responsibilities for students to help foster their site-level engagement and cohort building.

The example of the committee structure and the problems that students attributed to lack of structure highlight the overarching importance of structure. Indeed, a growing body of research indicates how structure in the form of policies and procedures helps to ensure equitable engagement and success of all students regardless of their backgrounds or prior preparation (Balster et al., 2010; Eddy & Hogan, 2014; Hurtado et al., 2008; Tanner, 2013). Science research itself is an unstructured or “ill-structured” endeavor, meaning that there are multiple ways to make progress and no single “right” answer (Dolan & Weaver, 2021; Simon, 1977). In addition, at least some of the students in this study struggled to organize their workdays because they did not have the structure of physically leaving home at a regular time to go to a research environment. Thus, remote research appeared to function as a “double whammy” – requiring students to navigate an ill-structured task in an unstructured environment. Students in sites that included more structure noted how this was a strength. In particular, students sought clear, consistent, and widely communicated schedules, expectations, and milestones as well as information about resources, such as who can provide help when issues or challenges arose. Students also wanted one-on-one meetings daily or every other day with mentors and meetings with their entire cohort and site leadership at least weekly. While some flexibility is needed and was expected, our results provide evidence that leaving structures entirely to individual research groups (e.g., whether and how frequently mentors meet with students) was problematic for students. Conducting an audit to identify technology needs in advance of the site start date is another example of a structure that would help to mitigate issues with diverse technology needs that students perceived as undermining their research progress and professional growth.

One of the most striking results in our view was how few students mentioned that they discussed issues related to diversity, equity, inclusion, or justice (DEIJ) during their programs. This result is especially noteworthy for multiple reasons. First, the NSF REU program prioritizes engagement of persons excluded because of ethnicity or race (Asai, 2020). Second, the sites took place just months after the killings of Ahmed Arbery, Breonna Taylor, and George Floyd and at the height of national consciousness about BlackLivesMatter, and all sites included one or more activities or events related to DEIJ. Furthermore, the #ShutDownAcademia / #ShutDownSTEM strike occurred on June 10, when all of the sites in this study were in session. It is possible that these discussions occurred and were simply not reported during focus groups. It is also possible that DEIJ activities or events were too limited in scope or disconnected from other aspects of site programming to be perceived as a strength. For instance, the one site where DEIJ was reported as needing improvement held multiple DEIJ events, but students perceived that people from excluded backgrounds were missing from non-DEIJ workshops or seminars. This finding brings to attention, once again, the need to restructure higher education such that DEIJ is an integral element rather than an additional activity.

Fortunately, there is a growing body of research on how to engage in difficult dialogues that can be used to ensure that REU sites dedicate time and create safe spaces for discussion of the value of diversity, ways to ensure equity and promote inclusion, and the importance of justice (Asai, 2020; Asai & Bauerle, 2016; Page, 2008; Sue et al., 2009; Tienda, 2013). At least some of this research has been described and translated into practical actions that could be applied to REU sites (Braun et al., 2018; Gin et al., 2020; Harrison & Tanner, 2018; Pfeifer et al., 2020; Seidel et al., 2015; Tanner & Allen, 2007; Tanner, 2013). Students at sites that created space and time for these discussions called them out as important conversations that helped them see their role in the world of science research. Future programming should ensure that time and space is dedicated to engaging in these important discussions and that the voices and experiences of people of color are integrated throughout programming, tapping local experts in diversity offices and centers for teaching and learning for guidance.

It is important to note that the study reported here is descriptive and evaluative in nature rather than a comparison of outcomes of remote versus in-person REU sites or a causal test of whether certain variables influence the effectiveness or inclusiveness of remote REUs. We have strived to keep our reporting of the results descriptive and, when possible, to highlight other research that is useful for understanding the observations and for improving remote REU sites in the future. Table 2 provides a list of the specific recommendations that students offered for maximizing the quality of their experience in remote REUs.

Our results raise several questions that should be addressed in future research. For example, what professional development and support structures are needed to ensure the quality and effectiveness of remote mentorship relationships? To what extent do remote REU sites allow engagement of undergraduates in research who would otherwise not have such opportunities? Do students in remote REU sites pursue graduate education and research related careers at the same level as students who complete in-person programs? Could REU sites that involve some students in person and others at a distance without creating inequitable experiences among members of the cohort or their mentors? Although these questions should be pursued with caution to avoid disadvantaging those who participate in research remotely, our results provide evidence that remote REUs are sufficiently positive to allow for further investigation of their affordances and constraints.

## ACKNOWLEDGEMENTS

We thank all of the students, faculty, and other research mentors for their willingness to proceed with remote REU programming and for sharing their experiences so that others could learn. We also thank Riley Hess for her feedback on drafts of this manuscript. This material is based upon work supported by National Science Foundation under Grant No. DBI-2030530. Any opinions, findings, conclusions, or recommendations expressed in this material are those of the authors and do not necessarily reflect the views of any of the funding organizations. The authors dedicate this work to all of the undergraduates seeking to do research and the individuals who provide these opportunities despite challenging circumstances.

---

**Figure 4.**
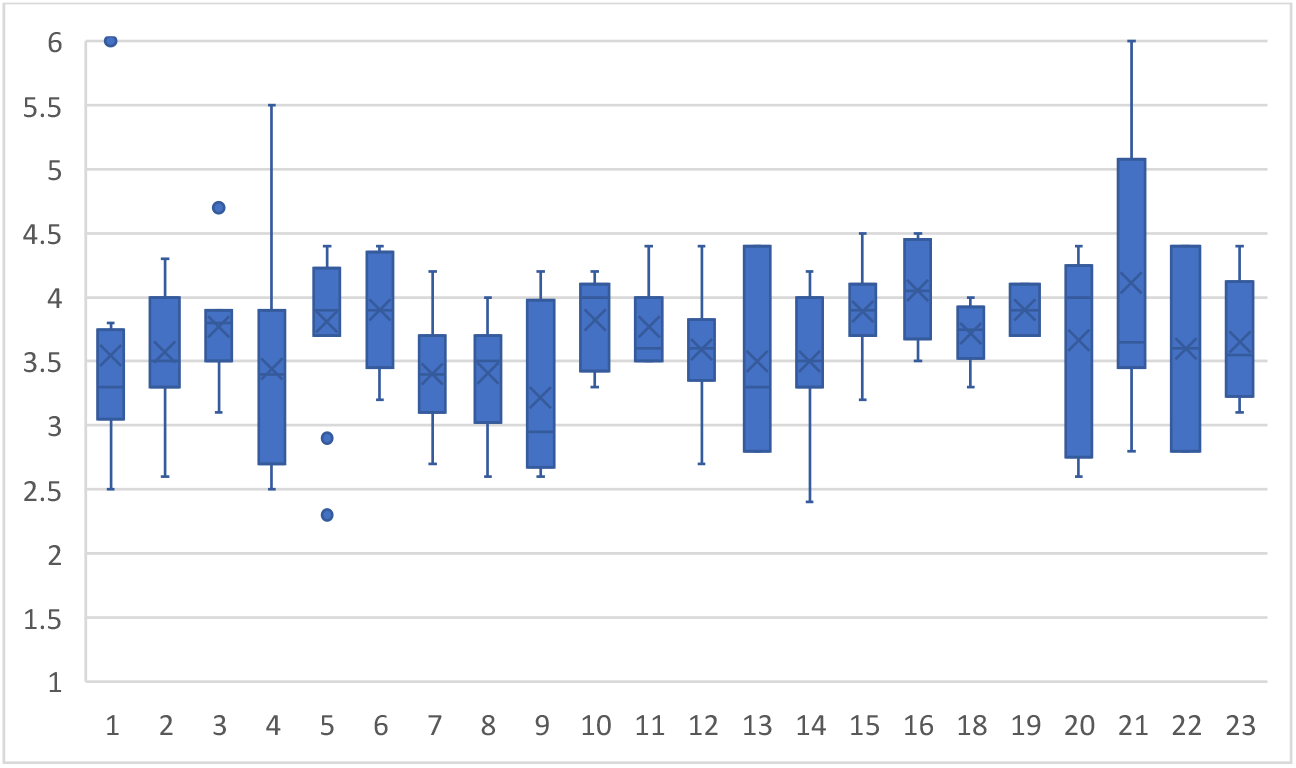
Connectedness. Students were generally positive about the sense of connectedness they felt in their program (*M*=3.6 out of 6; *SD*=0.6), but their ratings were lower (i.e., lower means and medians) and more consistent (i.e., smaller standard deviations) within each site than ratings of their relationships with their mentors. This figure shows student ratings by site, with 6 indicating strong agreement and 1 indicating strong disagreement with a positive statement about connectedness within the program (see supplemental materials for items and rating scale). The X signifies the site mean and the bar indicates the site median. Only data from remote students are included here (i.e., no responses from in-person students in programs 22 and 23).

**Figure 5.** Recommendations for Remote REU Sites. During the focus groups, students offered a number of recommendations for maximizing the quality of their experiences in remote REUs, compiled here.

## 10 STUDENT RECOMMENDATIONS TO MAXIMIZE THE QUALITY OF REMOTE REU EXPERIENCES

### 1 MENTORSHIP

- Ask how students are doing in general, not solely about their research experience. If comfortable, consider disclosing some information about how you are doing in general.
- If you are unable to help your student with a problem, connect them with someone who can.
- Establish open lines of communication early on (e.g., email, Slack, text) to ensure students feel comfortable reaching out with questions at times other than during regularly scheduled meetings.
- Ask for and listen to feedback from students about how you are doing as a mentor.
- Facilitate a balance between guiding students through their projects and allowing some autonomy to direct the research and answer their own questions.

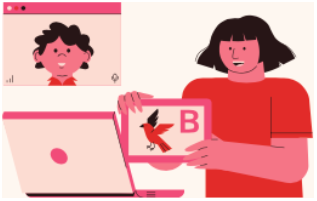

### 2 LEARNING

- Make explicit connections between what students are doing in their research and its relevance to “big picture” questions. For example, if a particular skill is used frequently in a field of a student’s interest, be sure to point out its utility.
- Give students opportunities to troubleshoot their problems on their own before providing answers or guidance.

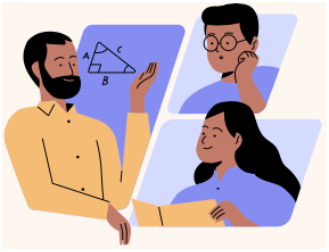

### 3 SENSE OF COMMUNITY

- Check with lab members to make sure they are willing to assist or provide guidance and then encourage collaboration within lab groups so that students feel comfortable going to anyone in the lab for help.
- Ensure that all students are aware of resources early on so they can make use of them if/when they need.
- Facilitate connections between students and other faculty or scientists in addition to their own mentors.

### 4 COHORT EXPERIENCE

- Utilize breakout rooms (or the equivalent) during meetings to give students opportunities to interact with one another.
- If possible, ensure the cohort is large enough for students to feel they are a part of something bigger than themselves.
- Hold regularly scheduled cohort meetings throughout the program, not only at the beginning.
- Facilitate informal interactions between students when possible. Consider holding synchronous, informal work time over Zoom to simulate an in-person work environment. Consider establishing a student-run Groupme or Slack for students to communicate with each other.
- Check in with students about how to structure virtual social activities to limit Zoom fatigue. Options include making these optional or holding them on days that no other meetings are scheduled.
- Facilitate open conversations on topics outside of research, such as current events, representation and DEIJ, and students’ roles in the scientific community.

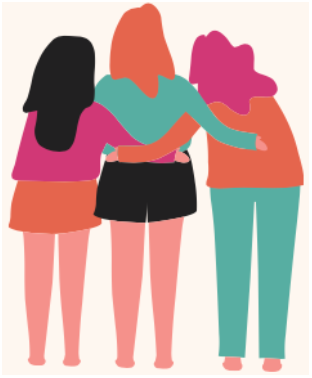

### 5 STRUCTURE

- Provide a daily or weekly schedule or a suggested schedule for students, including the number of hours of work expected per day and recommended breaks to prevent burnout.
- Help students make and recognize their progress by holding check-in meetings, establishing midpoint assignments, or setting intermediate goals.
- Distribute workload evenly throughout the program.
- Schedule skill-building sessions at a time in the program when students will be able to apply what they are learning.
- Ensure mentors set aside time every day or every other day to meet with mentees.
- Provide optional tasks or recommendations for what students could do if they have extra time.
- Host one or two sessions with mentors or program leadership to share and brainstorm strategies for time management (e.g., what to do, in what order, and when to get things done by).

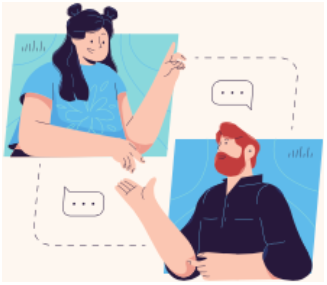

### 6 PROGRAM LOGISTICS

- Hold weekly program meetings to help establish connections and facilitate open communication.
- Be mindful of program pace. Keep consistent or slowly build up to ensure students are able to stay on track.
- Provide advanced notice of important dates and deadlines to help students gauge where they should be with their research and to give students and mentors sufficient time to plan.
- Limit the number of platforms to ease the logistics of communication.
- Ensure program leadership and mentors coordinate plans to minimize conflicts between programming and research.
- Stay within the confines of the original schedule as much as possible and minimize the number of unscheduled meetings.
- Break up lengthy online meetings to minimize Zoom fatigue.

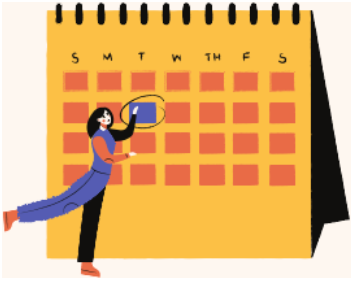

### 7 PROFESSIONAL SOCIALIZATION

- Provide opportunities for students to hear from current graduate students about their experiences in graduate school.
- Host sessions and panels highlighting a variety of research careers and the diversity of the scientific community.
- Provide information on the graduate application process and the myriad paths to graduate school.

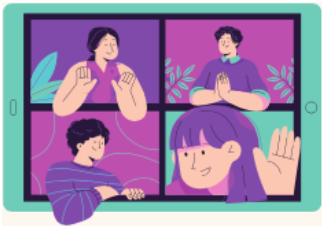

### 8 NETWORKING

- Ensure students have ample opportunities to meet, interact, and form relationships with faculty members, graduate students, and other members of the scientific community.
- Encourage students to collaborate with mentors and peers outside of their own lab group.
- Organize cross-site activities and events.

### 9 TECHNOLOGICAL ISSUES

Provide all students with the necessary login credentials and access information prior to program start.

Ensure in advance that students are supplied with necessary technology such as adequate computing capacity and reliable internet access.

In computation-focused programs, such as those that involve coding, be sure to provide resources and computation-specific support to students early in the program.

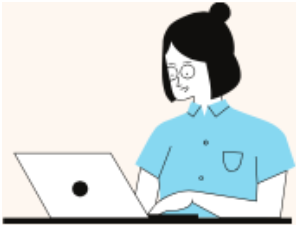

### 10 DIVERSITY, EQUITY, INCLUSION, JUSTICE AND REPRESENTATION

Provide repeated, formal and informal opportunities to discuss diversity, equity, inclusion, and social justice.

Ensure that all aspects of programs and programming include representation of individuals from backgrounds that are traditionally excluded or marginalized from the sciences.

Provide opportunities for students to hear from a wide variety of scientists and graduate school students who come from diverse backgrounds.

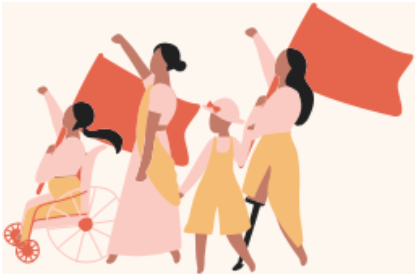

### RECOMMENDATIONS BASED ON

Erickson et al. “How do we do this at a distance?!” A descriptive study of remote undergraduate research programs during COVID-19

## Footnotes

1 Although this was not a focus of any of the discussions, it is important to note that the GRE is increasingly being dropped as a requirement for graduate applications in the life sciences and is not allowed to be reported by some programs. These decisions are driven by the growing number of studies showing the lack of predictive validity of the GREs for success in life science doctoral programs (e.g., Hall et al., 2017; Moneta-Koehler et al., 2017; see https://beyondthegre.org/grexit/ for a comprehensive list).

## Notes

### Competing Interest Statement

The authors have declared no competing interest.

