## Supplemental materials for "“How do we do this at a distance?!” A descriptive study of remote undergraduate research programs during COVID-19"

This supplement contains the following:


| Item | Page |
|---|---|
| Remote REU Site Profile Template | S2 |
| Remote REU Site Profiles (provided in alphabetical order by program name) | S4 |
| Focus group questions | S27 |
| Survey measures | S28 |

**REMOTE REU SITE PROFILE TEMPLATE**

**Scientific theme(s):**

- Program name:
- Link to your REU site webpage if you have one:
- Link to the public abstract for your REU site from the NSF website (or other funding source if applicable)
- Briefly describe the discipline(s) or general scientific themes of REU site
- Please indicate an author of the profile:

**Nature of the research:**

- Briefly describe the nature of the research that *typically* occurs in this site (bench, field, computational, other) in non-COVID years (if applicable)
- Briefly describe the nature of the research that occurred this year (bench, field, computational, other) and the extent to which the nature of the research had to change due to COVID-19.
- Were any new mentors recruited to the REU site because of the COVID-19 transition? For instance, were any mentors recruited this year because they had computational projects that were suitable for undergraduates? If yes, please briefly explain.
- Did experienced REU mentors have to generate / design new projects or research tasks because of the COVID-19 transition? For instance, were any mentors expecting to do field or benchwork who then developed a new/different project to make it possible to involve an undergraduate at a distance? If yes, please briefly explain.

**Program activities and expectations**

- **Time commitments:**
    - o About how many hours and/or days per week were students expected to work?
    - o To what extent were time commitments flexible or set?
    - o To what extent did the program set time commitments or let labs/mentors set time commitments?
    - o Briefly explain whether time commitments were handled differently because of COVID-19.
- **Structure:**
    - o Did mentors and/or or students work together differently than during previous summers because of the remote nature of the site? For instance, did students work in pairs or teams this year but not previously? Did mentors work collaboratively to mentor students this year but not previously? If yes, please describe.
    - o Please describe a typical week in the program to give a sense of how the program operated on a day-to-day and week-to-week basis.
- **Programming:**
    - o Please describe the program activities other than the research experience. Include brief descriptions of any formal (scheduled) aspects of the program, such as preparation to help students work remotely, training, professional development, workshops, panel

discussions, discussions of antiracism or social justice-related topics, social events, game nights, office hours, journal clubs, poster/talk presentations, etc.

- As much as possible, please indicate how **often** even type of event occurred (daily, weekly, monthly, once during the program), how **long** the event(s) typically lasted, and the context for the event / whether it was synchronous or asynchronous (e.g., synchronous zoom meeting). For example: “The program ended with a two-hour poster synchronous symposium on zoom during which each student presented their work for 10 minutes followed by 5-minutes of Q&A.”
- To the extent you know, did any informal (unscheduled or unstructured) activities occur related to the program? If so, please describe.
- Please describe any program activities intended for **mentors**, such as professional development on mentoring, discussions to plan remote-friendly projects, preparing mentors to work with students remotely, etc. Again, please include details about frequency and duration if possible.
- I**f applicable**, please describe any **new elements** of the program that you will continue to do in the future, including a **brief explanation of your reasoning**.

Please provide any other information that would be helpful for understanding how your REU site was designed and implemented, especially any differences or unique elements due to COVID-19.

### Biological Interactions Summer Research Program

PIs: Janet Branchaw & Amber Smith; General scientific theme: Interactions of Genotype and the Environment

Website: https://wiscience.wisc.edu/IBS-SRP Abstract: https://www.nsf.gov/awardsearch/showAward?AWD_ID=1659159

#### Programming

- The nature of research was computational.
- 12 students and 11 mentors
- 21-40 hrs/week expected depending on level of engagement; students either engaged in part-time research or full-time research
- Synchronous and asynchronous components

#### Mentors

- The mentoring structure mostly stayed the same as in previous years, but this year grad students and postdocs, "Badger Buddies," were also recruited to provide mentorship and support to students.

#### Influence of Pandemic

- The nature of research is typically bench and field research, but these kinds of research could not occur due to the pandemic.
- New mentors were not recruited as a result of the pandemic; students were already placed with mentors when the program shifted to a virtual format.
- All mentors had to redesign their projects to focus more on computation and experimental design so students could participate remotely.
- Students did not work in groups as a result of the pandemic.

#### Activities

Program Start

Program orientation

Graduate School Virtual Fair

Graduate School Admissions Panel

Bioscience Career Panel

Final Symposium

Program End

Throughout: Biological Interactions Seminars, Professional development seminars BI Drop-In Hours, Bioethics Book Group

| Activity Type | Description |
|---|---|
| Starting event | **Program orientation:** Much of the orientation was devoted to a series of community building exercises to help students get to know one another and practice using Zoom features to facilitate online engagement. The orientation also included information about program logistics, expectations, and how to access materials needed for the program. The session ended with a reflection on their readiness for research and a goals setting exercise. |
| Professional development | Weekly **professional development seminars** were held for students to gain skills and knowledge in navigating the research environment, develop effective mentor-mentee relationships, and explore psychosocial constructs (e.g., self-efficacy, imposter syndrome, growth mindset, stereotype threat), research careers, ethics, and diversity in STEM and science communication. **Graduate School Virtual Fair:** This event was an all-day fair in which students could come and go freely. Students had the opportunity to learn about graduate programs and connect with program staff to ask questions. **Graduate School Admissions Panel:** Graduate program directors and coordinators discussed the application process and qualities that make for a strong candidate. Students had the opportunity to ask questions. This event was one hour in duration. |
| Scientific | **Bioethics Book Group:** Students and Badger Buddies were invited to read *The Immortal Life of Henrietta Lacks* by Rebecca Skloott and discuss in a weekly book group. This summer a group of 8 students and Badger Buddies participated. This was an optional activity. |
| Research | **Biological Interactions Seminars:** Students met weekly to engage in activities and discussions about research and science careers. These sessions provided the opportunity to hone students' skills of observation, critical thinking, and creativity as they worked in small groups with a faculty expert to explore a phenotype. |
| Other | **BI Drop-In Hours:** Each week students were invited to join their peers and Badger Buddy mentors to ask questions about the program or questions about grad school, research, or career paths. Each week had a topic for discussion, but students could ask any question at any time. Social activities rounded out the hour if there was time after the discussion. |
| Ending event | **Final Symposium:** Students created graphical abstracts about their projects and gave a 5-minute presentation about their work over the summer and answered questions for 5 mins. Mentors and peers attended and asked questions. |

### Biological Research in Ecological and Evolutionary Developmental Biology BREED

PI: Cynthia Sarah Cohen; General scientific theme: Evolution, Development, & Ecology

Abstract: https://www.nsf.gov/awardsearch/showAward?AWD_ID=1659175

#### Programming

- Nature of research was computational, coding, and data analysis.
- 12 students and 4 mentors
- 40 hrs/week expected
- All activities were synchronous; student work was completed remotely.

#### Mentors

- There was some mentor collaboration/substitution due to a paternity leave, but this was not a result of the pandemic.

#### Influence of Pandemic

- The nature of the research changed from bench, field, and computational, to solely computational as a result of the pandemic.
- Mentors who had the ability, interest, and time were recruited; some were current, but two were new.
- All mentors had to redesign their projects to accommodate the remote circumstances.
- Some students worked in larger teams of 4-5 due to having fewer mentors as a result of the pandemic. Some students did work alone or in teams of 2 as well.

#### Activities

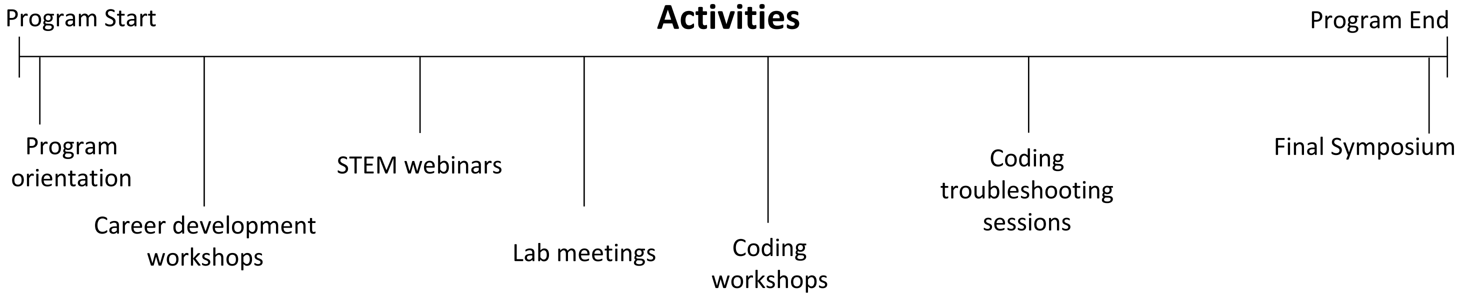


| Activity Type | Description |
|---|---|
| Starting event | **Program Orientation:** Participants met via Zoom for 2 hours, introduced themselves, and learned about program logistics and expectations from the program lead. Trouble-shooting regarding connectivity issues was carried out before the meeting. There was also a separate orientation for the SCIP coding workshop portion. Students from the same REU mentor lab group were mixed into several different coding groups to increase interactions across labs. |
| Professional development | **Career development workshops:** 3-hour long workshops were held via Zoom once a week jointly with other SFSU summer interns. Topics included discussion of professional development, working remotely, discussion of current events, ethics, antiracism, social-justice, writing an abstract, and presenting a talk. Sessions included why become a scientist, why go to graduate school, the PhD application process, lab practices, reading scientific papers, designing an experiment, building resiliency to stereotype threat, strategies for giving an effective talk, meetings with SFSU alumni in PhD programs, an abstract writing workshop, scientific career panel with invited speakers from diverse careers, and a science communication workshop. |
| Scientific | **Coding workshops:** Zoom workshops were held 4 days per week in the mornings. Students worked in teams, and the events were hosted by the SCIP program. Additional weekly SCIP webinars featured guest researchers, frequently from historically underrepresented groups in STEM. The guests discussed their path in science and answered questions. Speakers also discussed incorporating coding into their work as well as the challenges they encountered while learning to code. **STEM webinars:** These 1-hour presentations were typically held once a week. These webinars provided exposure to varied content related to ecology, field work, climate change, research seminars, and other virtual REU programs. Presenters included NGOs, conservation organizations, and researchers from RO1 and comprehensive universities. |
| Research | **Lab meetings:** Each participating mentor held weekly lab group meetings. Lab groups had additional scheduled and ad hoc sessions as needed. Some labs held REU intern specific weekly meetings as well. |
| Other | **Coding troubleshooting sessions:** Students in one group met with program leadership on occasion to troubleshoot coding problems they encountered. |
| Ending event | **Final Symposium:** Students gave 10-minute presentations on Zoom followed by a 3-minute Q&A session. This was part of an annual college-wide summer intern symposium with participants from many different programs and their guests including family, friends, and lab mates. Sessions ran concurrently with talks clustered by topics rather than programs. |

### Bruins-in-Genomics (BIG) Summer Undergraduate Research Program

PIs: Alexander Hoffmann & Hilary Coller; General scientific theme: Bioinformatics & Computational Biology

Website: https://qcb.ucla.edu/big-summer/

#### Programming

- The nature of research was computational.
- 12 students and 15 mentors
- ~40 hrs/week expected
- Synchronous and asynchronous components

#### Mentors

- There was one assigned day-to-day mentor who was directly responsible for their students, but often several lab members participated in mentoring. This was not as a result of the pandemic.

#### Influence of Pandemic

- Nature of research did not differ substantially from previous years; it was still computational.
- New mentors were recruited to the site because some faculty were less enthusiastic about mentoring additional students given their other responsibilities at work or at home during the pandemic.
- Experienced mentors did not have to redesign their projects, but they had to think more carefully, and re-work their mentoring styles and mechanisms to engage and guide students.
- Most, but not all, students worked in pairs this year.

#### Activities

Program Start

Pre-program preparation

Collaboratory workshops

Mentor/Mentee Check-in Surveys

Professional development workshops

BIG Summer Alumni Panel

Virtual tour of UCLA

Discussion

Teatime with faculty

Research Symposium & Awards Ceremony

Program End

Throughout: QCBio research seminars, Weekend Activities/Competitions, Office Hours, Teatimes with graduate students

| Activity Type | Description |
|---|---|
| Starting event | **Pre-program preparation:** A list of resources (videos) was sent to students prior to the start of the program to help familiarize students with relevant skills/concepts. |
| Professional development | **Professional development workshops:** These workshops were optional and held on weeks 4-8. They were 30 minutes to an hour in duration and were held via Zoom. Students attended relevant workshops on topics ranging from How to Create an Online Presence to How to Prepare a Poster Presentation. **BIG Summer Alumni Panel:** This optional one-hour event was held synchronously via Zoom. Students participated in a panel discussion with BIG SUMMER Alumni from 2015-2019. Topics of discussion included mental health in graduate school, how to select and apply to graduate school programs, and industry vs. academia. **Discussion:** This event was an optional synchronous talk with HBCU African -American, female students (led by Tracy Johnson) via Zoom. **Bioinformatics teatime with graduate students:** These were biweekly optional events in which students could meet with graduate students via Zoom. |
| Scientific | **Weekend Activities/Competitions:** Students participated in a combination of team-based and individual activities. The one-time BIG Summer Hackathon was a 2-day team-based, synchronous activity that took place via Zoom, while the T-shirt Design and Scientific Prose competitions were individual, one-time, asynchronous activities that were recorded and shared via the BIG SUMMER Slack Channel and email. These were optional. |
| Research | **Collaboratory workshops:** These mandatory workshops were held on weeks 1-3, were an hour in duration, and were held daily via Zoom. Students attended scheduled workshops relevant to their research projects assigned by their mentors. **QCBio research seminars:** These 30-minute seminars were held every week via Zoom. Students were exposed to cutting edge research presented by UCLA postdocs and faculty. |
| Other | **Mentor/Mentee Check-in Surveys:** These were given asynchronously during weeks 1, 2, and 4. Surveys were used as a means of checking in with both mentors and mentees and identifying any pressing issues throughout the program via email. **Virtual Tour of UCLA:** This was a one-hour, optional asynchronous event. **Office hours:** Optional daily, open office hours, synchronous via Zoom. Students scheduled appointments with the program manager to discuss progress in labs, projects, and admissions. **Teatime with Faculty:** During week 8, students had the opportunity to meet with select faculty to ask questions and receive guidance on admissions for UCLA graduate school programs. |
| Ending event | **Research Symposium & Awards Ceremony:** The program ended with a two-hour, synchronous theme-based Research Symposium followed by a 45-minute Awards Ceremony via Zoom. Students were allotted 5 minutes to present their research projects followed by a 10-minute group discussion and Q&A. Students and faculty were recognized for mentorship and research excellence at the Awards Ceremony. |

#### Cary Institute of Ecosystem Studies REU

PI: Alan Berkowitz and Kevin Burgio; General scientific theme: Translational Ecology

Website: https://www.caryinstitute.org/eco-inquiry/reu-program Abstract: https://www.nsf.gov/awardsearch/showAward?AWD_ID=1559769

##### Programming

- Nature of research was translational ecology.
- 12 students and 20 mentors
- 35-40 hrs/week expected
- Synchronous and asynchronous components

##### Mentors

- Several projects already had a team of mentors (either several senior scientists, or a senior scientist and post doc or grad student) which allowed for increased student support.

##### Influence of Pandemic

- The nature of research was shifted to focus more on computation rather than field work and data collection.
- The same pool of mentors was maintained despite drastic changes to the program.
- Most of the mentors involved had to alter the subjects of student projects to accommodate for the altered nature of research.
- There were four research teams, each comprised of 3 REU students and a Cary post doc to provide additional peer-to-peer support and scientist-student mentoring.

##### Activities

Program Start

- Developing Research Questions and Hypothesis
- Research Proposals
- Data Inquiry Strand
- Scientific Writing and Communication workshops
- Translational Ecology (TE) in Action Panels
- Anti-racism and Social Justice discussion
- Writing a CV
- Career Pathways
- Creating a "Data Nugget"
- Giving Presentations workshop
- Personal Statement on TE
- Cary Undergraduate Research Symposium

Program End

Throughout: Research team meetings, Weekly "Tea Times," Check-ins with the program coordinator, student-organized social nights

| Activity Type | Description |
|---|---|
| **Starting event** | Prior to program start, students interacted with mentors, read papers, and looked at data. **Welcome, 50 Questions, My Place Profile**: 6 hours of synchronous and asynchronous work in week 1, introducing program opportunities, expectations, and norms in order to build cohort cohesion. **Mentor/mentee contract** completed in week 1. |
| **Professional development** | **Science communication:** 5 one-hour synchronous workshops with individual assignments led by Cary Institute's communications staff**. Creating a "Data Nugget":** 2 one-hour synchronous workshops led by Cary's education staff in basic strategies for ecology education were applied to developing a draft Data Nugget. **Career Pathways:** a one-hour synchronous workshop led by one of Cary's scientists and 2 postdocs. **TE in Action Panels:** 4 one-hour synchronous panel discussions, led by 2-3 people in various non-academic sectors on the topics of Science Communication, Ecology in the private sector, Environmental advocacy, and Ecology in NGOs. Students responded to reflective prompts before and after each panel. **Personal Statement on Translational Ecology:** Students completed a statement presenting their perspectives and expertise in Translational Ecology suitable for use in career and graduate school applications. **Writing a CV and contacting potential grad school advisors:** A one-hour synchronous workshop led by the REU program coordinator. |
| **Scientific** | **Data Inquiry strand:** Data carpentry and GitHub Carpentry online modules, 6 one-hour synchronous R workshops, and 2 one-hour synchronous Metadata workshops. **Literature searches** and **using citation managers:** 3 one-hour synchronous workshops led by Cary Institute's librarian. **Writing workshops:** 4 one-hour synchronous sessions led by the REU program co-director and coordinator. **Giving presentations:** A one-hour synchronous workshop led by the REU program coordinator. |
| **Research** | Students conducted independent research projects of their own design. **Research Proposals:** Students submitted 5-page proposals in week 3 that were reviewed by mentors, another Cary scientist, and a fellow REU student. **Research Teams** comprised of 3 REU students and a Cary Postdoc who provided additional mentoring and support. **Research paper:** Students chose to write a basic final report, a paper to be 'published' by Cary in its on-line Undergraduate Research Reports collection, or a paper to be submitted to a peer-reviewed journal. |
| **Other** | **Weekly "Tea Times":** 7 one-hour informal synchronous discussions with scientists from various ecology fields to discuss different aspects of being a scientist. **Anti-racism and social justice discussion:** A one-hour synchronous discussion led by the REU program coordinator. |
| **Ending event** | **Cary Undergraduate Research Symposium:** 7-minute presentations with 3 minutes for questions. **Focus group reflections**: 2-hour discussion of advice for future REU students, mentors and program leaders. |

#### Exploring 21st Century Careers in the Biological Sciences: A Comparative Regenerative Biology Approach

PI: Jane E. Disney; General scientific theme: Comparative Regenerative Biology and Aging
Website: https://mdibl.org/education/undergraduate-opportunities/reu/
Abstract: https://www.nsf.gov/awardsearch/showAward?AWD_ID=1851962

#### Programming

- The nature of research was computational, related to regenerative biology and aging or COVID-19.
- 6 students and 5 mentors
- 40 hrs/week expected
- Synchronous and asynchronous components

#### Mentors

- Mentors and/or lab group members met students 1:1 multiple times per week.
- Two faculty members co-mentored a pair of students; another faculty member transitioned a pair of students to Covid-19 projects when datasets became available.

#### Influence of Pandemic

- The nature of research typically includes benchwork components, and this year it was changed to be solely bioinformatics research.
- New mentors were not recruited as a result of the pandemic.
- All research projects were changed to accommodate the need for students to work remotely.
- Two students worked together on two separate bioinformatics projects as a result of the pandemic.
- The director of the Bioinformatics Core at MDIBL created a Bioinformatics Training Program to support students and mentors during this time.

#### Activities

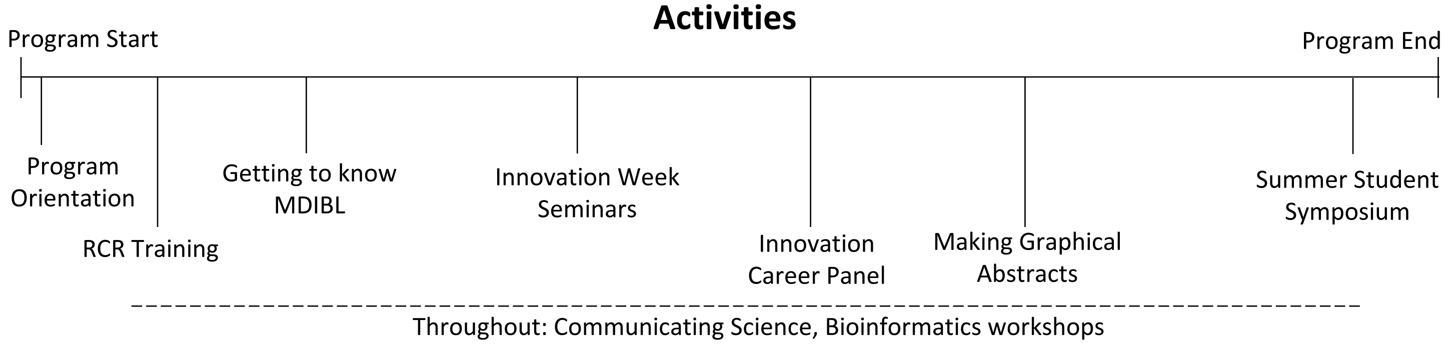


| Activity Type | Description |
| --- | --- |
| **Starting event** | **Program orientation:** Students were given a PowerPoint presentation by the president of the MDI Biological Laboratory on the history and current research of MDIBL. Representatives from the Finance, Education, Human Resources, and Development departments were present and reviewed campus processes and procedures with students. Students reviewed schedules of upcoming seminars, workshops, training opportunities, and other activities for the summer and had opportunities to ask questions. |
| **Professional development** | **Communicating Science:** All REU research fellows participated in a weekly course called Communicating Science. Students were introduced to best practices in quantitative communication, scientific writing, and in delivering formal scientific talks. **Bioinformatics workshops:** In a 2-part Bioinformatics introduction and follow-up weekly workshops, students gained an understanding of the importance of viewing computational and statistical analysis as an integral part of their experimental work. Aspects of this training included statistical assessment of projected experiments before data were collected. **Innovation Career Panel:** Students engaged in a discussion with a panel of innovative entrepreneurs. Each panelist introduced themselves, explained where they are today, and addressed the concept of innovation thinking and how they have applied it to their work. |
| **Scientific** | **Innovation Week seminars:** Students participated in 3 one-hour seminars, Innovation Thinking: An Introduction to Innovation Skills, Technology Transfer and Intellectual Property, and Case Studies from the Innovation Cohort at Maine Medical Center as a part of their Innovation Week activities. |
| **Research** | **Responsible conduct of research (RCR) training:** All students participated in a 2-hour RCR training given by the program's PI, Dr. Jane Disney. It included case studies for students to review and discuss. Students also participated in role playing scenarios in Zoom breakout rooms where they enacted events that might occur between students, mentors, and human resources. |
| **Other** | **Getting to know MDI Biological Laboratory:** Students viewed a "live" virtual tour of the campus and laboratory spaces and participated in a meet-the-faculty Zoom session, where each member of the research faculty introduced themselves to the students and described their research programs and invited questions from students. |
| **Ending event** | **Summer Student Symposium:** The culminating event of the program involved three days of sharing research. The first two days were asynchronous and were dedicated to students' Poster Sessions in which each student uploaded a biography, abstract, and poster. Participants could leave comments and questions for students to address. The last day was dedicated to students' oral presentations in the format of a Three-Minute Thesis. |

### Fungal Genomics and Computational Biology Summer Research Program

PIs: Jonathan Arnold; General scientific theme: Fungal Genomics and Computational Biology
Website: https://www.genetics.uga.edu/fungal-genomics-and-computational-biology
Abstract: https://www.nsf.gov/awardsearch/showAward?AWD_ID=1946937

#### Programming

- The nature of research was genomics and computational biology.
- 13 students and 11 mentors
- 40 hrs/week expected
- Synchronous and asynchronous components

#### Mentors

- There was one collaborative project on COVID-19 in which mentors worked together, but each REU participant had their own project within the larger project.

#### Influence of Pandemic

- Nature of research was shifted to focus solely on mathematical sciences and work that could be done on the computer.
- More computational biologists were recruited to the site to replace those mentors that put students on the bench.
- Some bench scientists designed new projects that could be done computationally.
- Some students did work collaboratively as a result of the pandemic; however, each REU participant had their own research project.

#### Activities

Program Start

Program End

Computer training

George Floyd Discussion

Responsible Conduct of Research Seminars

Final Symposium

Throughout: Weekly student-organized luncheons, research seminars, Zoom meeting with other REU partners

| Activity Type | Description |
|---|---|
| Starting event | **Computer training:** REU participants participated in a computer training throughout the first week of the program. |
| Professional development | **Bioethics/Responsible Conduct of Research Seminars**: These seminars were held once a week by UGA. Topics of these seminars included: Bioethics of CRISPR, Best Lab Practices and Academic Freedom, Science Communication, Risk Communication, Use of Sex and Race Categories in Research, Mentoring, and Collaboration and Workplace Rights. |
| Scientific | **Research Seminars:** These seminars were held every Thursday and dealt with various topics including a COVID19 research seminar. |
| Research | **Luncheon meetings with mentors:** Students met with REU mentors every week on Tuesdays.<br><br>Mentors met with REU participants 1 to 3 times per week via Zoom. |
| Other | **Student Luncheon:** Students self-organized weekly luncheons. **Zoom meetings with other REU partners. Luncheon meetings with mentors:** Students met with REU mentors every week on Tuesdays. |
| Ending event | **Final Symposium:** Students, gave a 10-minute presentation about their work over the summer, and answered questions for 5 mins. Mentors and peers attended and asked questions. These presentations were then uploaded to YouTube. |

### Genes & the Environment: Research Experiences for Undergraduates from Rural & Tribal Colleges at the University of North Dakota (UND)

PI: Van A. Doze; General scientific themes: Molecular & Cell Biology, Epigenetics, Gene Regulation, Computational and Systems Biology, & Neuroscience.

Website: http://ndinbre.med.und.edu/NSF-REU/ Abstract: https://www.nsf.gov/awardsearch/showAward?AWD_ID=1852459

#### Programming

- The nature of research was computational and systems biology.
- 25 students and 11 mentors
- 40 hrs/week expected
- Hybrid program with some (7) students participating in in-person activities and most (18) completely remote.

#### Mentors

- Mentors did not work in teams or collaborate more as a result of the remote format.

#### Influence of Pandemic

- Nature of research did not differ substantially from previous years.
- No new mentors were recruited as a result of the pandemic, although only 11 of the 20 original mentors decided to participate.
- Two mentors developed entirely new projects due to the remote format, and others had students work on aspects/tasks of the lab's research projects which could be done remotely (e.g., data analysis, genomics).
- Some labs that had less experienced students had students work in groups.

#### Activities

Program Start

Five-in-five

Women in STEM Panel

Pub Med Searching

Techniques: Electrophoresis & Immunoblotting

Stressors Discussion

Difficult Conversations

Stressors Discussion

Mindfulness Practices Now

Grad School Showcase & Prep Panel

Final Research Symposium

Program End

Throughout: Bioethics seminars, UND faculty seminars, UGA faculty seminars, Departmental seminars, meetings with mentors, journal clubs, lab meetings

| Activity Type | Description |
|---|---|
| Starting event | **Five-in-five:** Each student presented 5 slides about themselves as an introduction; several faculty/staff participated. This event continued for approximately 3 weeks with 1-2 meetings/week. It was held synchronously via Zoom and was 1-2 hours in duration. |
| Professional development | **Bioethics/Responsible Conduct of Research Seminars**: These seminars were held once a week by UGA. Topics of these seminars included: Bioethics of CRISPR, Best Lab Practices and Academic Freedom, Science Communication, Risk Communication, Use of Sex and Race Categories in Research, Mentoring, and Collaboration and Workplace Rights. **Virtual Grad School Showcase and Grad School Prep Panel. NSF-GRFP Presentation Graduate Research Fellowships Program. Women in STEM – Making Smart Choices for Your Career Panel Discussion:** This synchronous presentation was facilitated by 3 UND female science faculty members. |
| Scientific | **UND faculty research seminars:** These seminars were held synchronously once a week via Zoom and lasted 1-1.5 hours in duration. **UGA faculty research seminars:** These seminars were held synchronously once a week. They included individuals in a wide variety of research fields who shared their academic/career journeys, many of which were not traditional. **Departmental seminars:** UND graduate students gave seminars and/or defended their thesis. These seminars were optional, held synchronously via Zoom, and were one hour long. |
| Research | **Pub Med Searching:** This event was held synchronously via Zoom, was one-hour in duration, and was provided by UNDSMHS Librarian. **Techniques: Electrophoresis and Immunoblotting:** This presentation was held synchronously over Zoom and was given by a Biomedical Sciences faculty member. |
| Other | **Meetings with mentors, journal clubs, lab meetings:** These varied by lab. **Stressors Discussion Sessions (COVID19 stress; racism; riots):** Two Zoom sessions facilitated by UNDSMHS Assistant Director of Interprofessional Education. **Difficult Conversations:** One synchronous Zoom session facilitated by UNDSMHS Assistant Director of Interprofessional Education. **Mindfulness Practices Now:** One synchronous/Zoom session: An exploration of mindfulness practices, evidence-based benefits, and how you can begin to apply mindfulness to your life to manage your body's stress response. |
| Ending event | **Final Research Symposium:** The program concluded with an all-day synchronous symposium on Zoom. Each student presented (on PowerPoint) their work for 10 minutes with 5 additional minutes for questions. There were 3 groups of 2-3 students; all others were individual presentations. |

### iCompBio at the University of Tennessee at Chattanooga

PI's: Hong Qin & Soubantika Palchoudhury; General scientific theme: Computational and Quantitative Biology

Website: https://www.utc.edu/faculty/hong-qin/icompbio.php

Abstract: https://www.nsf.gov/awardsearch/showAward?AWD_ID=1852042

#### Programming

- The nature of research was computational and quantitative biology.
- 40 hrs/week expected
- 12 students and 12 mentors
- Synchronous and asynchronous components

#### Mentors

- Mentors typically met with students two to three times during the week.
- PI Qin typically held joint meetings to ensure all participants were making progress on their research.

#### Influence of Pandemic

- Nature of research did not differ substantially compared to previous years.
- The iCompBio lab typically was designed for 10 undergraduates, however, this year that number was increased to 12 since research was virtual.
- Several faculty members joined the research team due to cancellations of other REU programs across campus.
- Approximately half the mentors proposed new research projects due to the pandemic.
- Each student had their own individual research project and mentor that they were assigned.

#### Activities

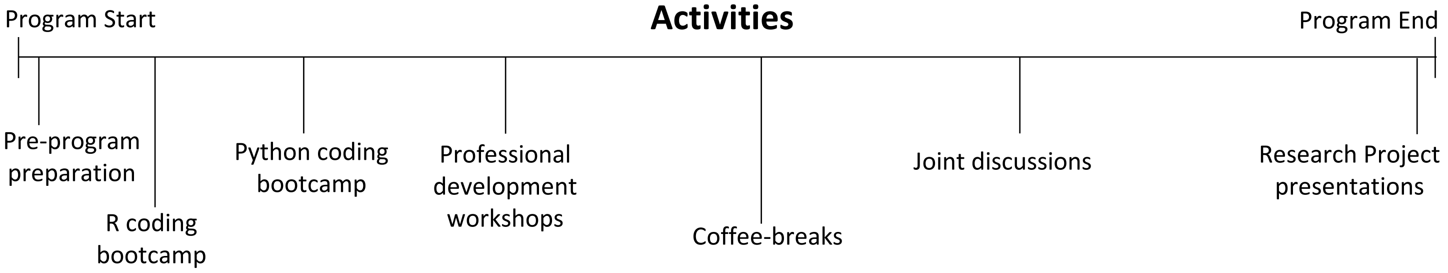


Throughout: Cohort meetings, mentor-mentee meetings, game nights, joint working sessions

| Activity Type | Description |
|---|---|
| Starting event | **Pre-program preparation**: This served as orientation and introduction to the program for student. Participants completed the CITI Responsible Conduct of Research program as well. |
| Professional development | The program hosted a formal **presentation training workshop** where they discussed the best practices of research presentations, followed by a working session on presentations in which students competed amongst one another to present a small presentation. Other students would then provide critical feedback for ways to improve. Students met with mentors on a frequent basis to discuss research. Additionally, all students attended an **NSF GRFP webinar.** |
| Scientific | Collaborative online documents were used to share reading notes and research progress with mentors and PIs. **R coding bootcamp:** This was a week-long program in which students went over basic data frame and computational programming skills such as retrieving and analyzing data. **Python coding bootcamp:** This was an intensive two-day programming bootcamp that focused on data analysis and data visualization. |
| Research | **Weekly group meetings:** Students discussed research progress and plans for the week. **Joint working sessions:** Students discussed their projects in groups to inform and receive advice from others on their individual projects. **Open virtual office hours – "coffee breaks":** Students were welcome to join a zoom session to talk about research and ask questions with PI Qin. |
| Other | **Two virtual game nights** were hosted and planned informally by students within the program as well as with students outside of the REU program. Faculty were also invited to take part in these activities. |
| Ending event | **Research Project Presentations:** During the last two days of the ten-week program, all students gave 25-minute presentations on their research project and then had 5 minutes for Q&A. Family, friends, and professors from other universities were also invited to watch student presentations. |

### Indigenous America to Indigenous Mekong – Adventures in Biology and Biodiversity

PI's: Ruben Michael Ceballos, Danielle Levesque, and Elizabeth Padilla-Crespo; General scientific theme: Biology, Microbiology, Biodiversity, and Genetics

Website: https://ceballoslab.uark.edu/nsf-reu-iaim/

#### Programming

- The nature of research was predominantly in silico with data analysis, applied bioinformatics, and computational biology.
- 40 hrs/week expected
- 14 students and 12 mentors
- Synchronous/Asynchronous components

#### Mentors

- Students and mentors were required to meet at least once every two days for one hour to discuss projects.
- Two "all-hands" meetings were scheduled once a week: one between all students and the PI, and the other with all students, mentors, and the PI.
- PI regularly had one-on-one meetings with project mentors.

#### Influence of Pandemic

- Nature of research differed from previous years, as this is typically an international REU program. International travel was suspended by the awardee institution causing most projects to be redesigned to be in silico, or computer-based.
- In previous years, students worked on team projects with each carrying out an independent component of the team project.
- Three additional mentors were added to the research team as a result of the pandemic. The mentor pool consisted of PIs from different universities whose programs had been impacted by pandemic, allowing them to join this program remotely making for a diverse set of students, mentors, and projects.
- The Plant Metabolomics Data Analytics REU Site (PI: Anne Osano, Bowie State University; https://www.nsf.gov/awardsearch/showAward?AWD_ID=1757607 ) joined with this site for a larger cohort and program experience.

#### Activities

Program Start — Program End

1. Virtual Introductory Sessions
2. Technical skills training
3. Professional development activities
4. Technical skills training
5. Professional development activities
6. Student Research Presentations

Throughout: Workday meetings, "all hands" meetings, mentor outreach, and weekly professional development activities.

| Activity Type | Description |
|---|---|
| Starting event | **Virtual introductory sessions-** This consisted of two introductory zoom meeting in which PI Ceballos set forth expectations for students and mentors on a daily basis. The first meeting acted as an introductory session and general overview of the program between PI Ceballos and student participants. The second zoom meeting included day to day mentors and laid forth general guidelines for the summer. |
| Professional development | **Weekly Professional Development activities:** These took place each Wednesday during the program and lasted anywhere from 1 to 3 hours. Theses included technical skills training over topics such as laboratory safety and research ethics. Additionally, these professional development activities included scientific methods and career training which differed on a group-by-group basis. Some of this training included learning how to work with NAMD and VMD software programs as well as bioinformatics packages. These activities also included talks on how to apply to graduate school. |
| Scientific | As mentioned previously, technical skills training during the first half of the program encompassed programing as well as safety and research ethics. |
| Research | **Workday Meetings**: Each morning, or every other morning, students had workdays meetings with their mentor to discuss what they would accomplish on their research that day. Additionally, the PI Ceballos would meet with all students on Fridays to discuss how projects were going as well as gauging mentor performance. |
| Other | **Mentor outreach**: Mentors were encouraged to contact students each day around 8-8:30 AM to ensure students were awake and working on their research projects. Additionally, students were encouraged to meet and collaborate amongst themselves outside of mentor engagement. |
| Ending event | **Student Research Presentations**: The program allocated two days at the end of the program to students' presentations on their research. These presentations were approximately 10 minutes each with an additional 5 minutes allocated to Q&A. |

### Integrative Biology and Ecology of Marine Organisms

PIs: Adam Summers & Stacy Farina; General scientific theme: Marine biology, ecology, development, physiology, and biomechanics
Website: https://fhl.uw.edu/research/summer-research-internships/
Abstract: https://www.nsf.gov/awardsearch/showAward?AWD_ID=1852096

#### Programming

- The nature of research was lab and field experience.
- 11 students and 11 mentors total; 2 students and their mentors worked remotely
- 30-40 hrs/week expected
- All hybrid with in-person field trips

#### Mentors

- There was one situation where an on-site mentor supplemented the mentoring of an off-site mentor.

#### Influence of Pandemic

- Nature of research did not differ substantially from previous years.
- No new mentors were recruited to the site as a result of the pandemic.
- Some mentors switched to more computationally intensive projects because the lab work was not possible.
- Students did not work in pairs or teams as a result of the pandemic.

#### Activities

Program Start

Virtual Orientation

Virtual Dinner

BLM, Antiracism, and Pride events

Trawl trip

Group tour of Salish Sea

Tidepooling trip

Final Presentations

Program End

Throughout: Friday Harbor Labs Seminars, Luncheon Seminars, Professional Development Workshops

| Activity Type | Description |
| --- | --- |
| **Starting event** | **Virtual Orientation:** Students traveled to the site two weeks early to quarantine. During these two weeks, students met outside and participated in various team-building activities. Additionally, students and mentors introduced themselves, and mentors gave short introductions about what they do at the site. **Virtual Dinner:** Everyone who was involved with the REU site in some manner participated in a virtual dinner event over Zoom. |
| **Professional development** | **Professional Development Workshops**: Students participated in weekly professional development workshops, which included a graduate student panel, a grant workshop, a mock interview workshop, a writing workshop, an experimental design and statistics workshop, a literature review workshop, a bioethics workshop, and RCR training. |
| **Scientific** | **Group tour of the Salish Sea:** Students engaged in a group tour of the Salish Sea which was meant to give students a general appreciation for the wildlife they were studying. **Trawl Trip:** The REU participants were taken out on a fishing trawler. Students learned how to deploy trawl gear, catch fish, and sort the fish. **Tidepooling Trip:** Students ventured to nearby tide pools to examine the fauna living within them. |
| **Research** | **Friday Harbor Labs Seminar Series:** Professors, graduate students, and/or post-docs that were doing research or had previously done research at Friday Harbor labs delivered presentations to students virtually. These were held weekly. **REU Luncheon Seminars:** Professors and researchers delivered 40–50-minute presentations to students, and students were encouraged to ask questions and participate in an open, informal discussion. |
| **Other** | **BLM, Antiracism, and Pride Events:** Students made posters and banners in honor of the Black Lives Matter movement and Pride month. Students were encouraged to reflect on current events and have conversations with one another about what was going on at the time. **Outdoor game activities:** Students participated in 2 game nights during the program. |
| **Ending event** | **Final Presentations:** At the beginning of the final week of the program, students delivered their presentations via Zoom. The whole island was invited to the symposium. Students were given feedback to incorporate into their final papers, which were due at the end of the final week. |

#### The Interdisciplinary and Quantitative Biology Research Experience for Undergraduates (IQ BIO REU) at the University of Puerto Rico

PI: Juan Ramirez Lugo & Patricia Ordoñez; General scientific theme: Bioinformatics, Genomics, Computational, and Quantitative Biology.

Website: http://iqbioreu.uprrp.edu Abstract: https://www.nsf.gov/awardsearch/showAward?AWD_ID=1852259

#### Programming

- The nature of research included quantitative and computational biology, gene expression analyses, bioinformatics, and land use dynamics in tropical ecosystems.
- 37.5 hrs/week expected
- 12 student and 7 mentors
- Synchronous/Asynchronous

#### Mentors

- Participants were recommended to meet with the PIs and mentors at least once a week to discuss research progress. PIs required "Monday Check-Ins" with students to ensure mentors and students were communicating frequently.

#### Influence of Pandemic

- Nature of research changed to being strictly computational which differed in comparison to past years where computational work was combined with field and laboratory work as well.
- No new mentors were recruited to the site, and fewer mentors participated in the program compared to past years since not all projects were able to transition remotely.
- In previous years, students worked alone in labs. However, this year, students were paired together in lab groups to allow for collaboration and social bonding in the new virtual format, but each student had their own individual research project.

#### Activities

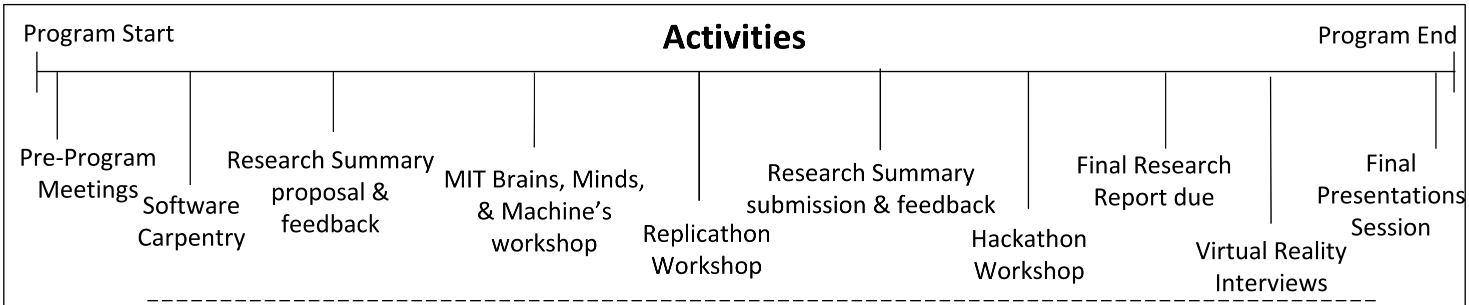


Throughout: Professional development sessions and Student meetings with PI's and mentor, workshops, and movie nights.

| Activity Type | Description |
|---|---|
| **Starting event** | **Pre-Program Meetings**: Prior to the program beginning, researchers held two remote meetings with students to help layout expectations for the summer and respond to any potential questions or concerns. These meetings were also intended to help promote social engagement and cohesion amongst students. |
| **Professional development** | **Professional Development Sessions:** During the first 5 weeks of the program, 2-4 professional development sessions were held each week for approximately 1-1.5 hours. During the last 4 weeks of the program these sessions occurred only 1-2 times a week as student feedback indicated participants wanted more time to focus on research. **Virtual Reality Interviews**: At the end of the program, students used a virtual platform, Mozilla Hubs, to discuss their research with other students. This session lasted for about one hour, during which students had very brief one-on-one discussions about their research projects using virtual avatars. |
| **Scientific** | **Software Carpentry:** These workshops took place during the first 7 days of the program and students spent ~4 hours each day in workshops learning basic coding skills, statistical research skills, and programming. During these workshops students were placed into Zoom breakout rooms to allow for collaboration of ideas.<br>**Replicathon Workshop:** This was a two-day workshop dedicated to data reproducibility using R studio. **MIT Center for Brains, Minds and Machines Workshop:** This was a two-day workshop focused on three core themes: computation, neuroscience, and cognition. **Hackathon Workshop:** Students would attempt to hack into their own projects through collaboration with peers and local mentors from all types of different backgrounds in hopes of improving or expanding upon their research. |
| **Research** | **Research Summary Submission & Feedback:** During both the 3rd and 6th week of the program, students submitted a research summary and received feedback. Students were expected to use feedback to then make improvements that would be incorporated into their final report. |
| **Other** | Students participated in the **#ShutdownSTEM** movement taking place across the country. During this time students discussed implicit bias, and this was followed by a movie addressing themes of social justice. **Movie Nights**: Each week following the groups participation in the #ShutdownSTEM movement, the group decided to have weekly movie nights which consisted of films that touched on topics of racial injustice and race relations. |
| **Ending event** | **Final Presentations session**: The program ended by hosting a day-long final presentations session. During these presentations, students gave 10-minute presentations and had 3 minutes for Q&A. |

### Mitigating the Impact of COVID-19 Pandemic on Undergraduate Research Training in the Biosciences, Microbiology & Immunology Dept

PIs: Margaret (Mari) Eggers & Colin Shaw; General scientific theme: Environmental Biosciences

Abstract: https://www.nsf.gov/awardsearch/showAward?AWD_ID=2034045

#### Programming

- The nature of research was Community-engaged Tribal-University partnerships addressing environmental bioscience issues of importance to Montana Tribes.
- 8 students and 9 mentors
- 40 hrs/week expected
- Hybrid program with students at Tribal colleges participating in in-person field work and all students interacting with mentors and completing research remotely.

#### Mentors

- Mentors worked directly with PIs to recruit students for the remote program.
- The entire cohort met virtually for one hour per week with PIs.

#### Influence of Pandemic

- This was a one-year program to increase student research opportunities during the pandemic, with both the Microbiology and Earth Sciences Departments starting regular REU programs in 2021. The nature of research was designed to offer a REU experience for students, especially Native American students, whose previous research plans were canceled due to the pandemic. The projects combined fieldwork in Tribal communities with lab work at MSU.
- Although the program was specifically designed with COVID-19 restrictions in mind, when reservations closed their borders mid-summer, the group was forced to make adaptations to the joint field work planned for Tribal and MSU members such as finding other local fieldwork opportunities. MSU based students gained more lab experience, while Tribal based students gained more field work experience.
- Students based at MSU had one primary mentor and support from their lab groups, and students at the Tribal colleges had both an MSU mentor and a mentor from their home community.

#### Activities

Program Start

Program End

Program orientation

Introduction to Research

CITI RCR Training Certificate

Virtual graduate school fair

Virtual Research poster presentations

Throughout: Weekly cohort meetings, professional development activities, GRE preparation, seminars, and social justice discussions

| Activity Type | Description |
|---|---|
| Starting event | **Program Orientation**: The first week of the program served as the orientation session for participating students. |
| Professional development | **GRE Preparation**: The program offered optional GRE preparation which was beneficial for many of the students already thinking about applying for graduate school. **Professional Development Activities**: All students were required to complete CITI RCR training during the program. Additionally, the program allowed for students to pick from an array of professional development activities to work on each week for a minimum of 8 hours. **Virtual graduate school fair**: During the $5^{th}$ week of the program students were given the opportunity to participate in the National Virtual Graduate School Fair for 2020 REU scholars. |
| Scientific | **Research and Ethics seminars**: Students attended numerous seminars from the NSF REU program throughout the duration of the program. |
| Research | **Introduction to research**: The second week of the program acted as an introduction to research for participating students. **Weekly Cohort meetings:** All participants in the program would meet virtually via Webex for one hour a week with the PI with a focus on both developing and creating a research poster to virtually present their project at the end of the program. During the first half of the summer, collaborative field-work took place and was dependent on each respective tribe's pandemic policies. |
| Other | |
| Ending event | **Virtual Research Poster Presentations**: At the end of the program students virtually presented their research posters. |

### RAPID: Undergraduate Research in Modeling and Computation for Discovery of Molecular Probes for SARS-CoV-2 Proteins

PI's: Mary Jo Ondrechen, Steven A. Lopez, and Mona Minkara; General scientific theme: Biochemistry, Computational Biology, Computational Chemistry

Website: http://nuvreu.appspot.com/

Abstract: https://www.nsf.gov/awardsearch/showAward?AWD_ID=2031778&HistoricalAwards=false

#### Programming

- The nature of research was computational.
- 8 students and 3 mentors
- 35 hrs/week expected
- Synchronous and asynchronous components

#### Mentors

- Mentors did not collaborate with one another or work in teams as a result of the pandemic.

#### Influence of Pandemic

- This was a one-year project designed specifically for the pandemic and funded under the RAPID mechanism.
- All three mentors joined the project because of the needs that arose from the constraints of the pandemic.
- The projects were designed specifically to be done remotely, but this does not represent a significant change in the operations of the three PIs.
- Students did not work in teams as a result of the pandemic.

#### Activities

Program Start

Program Orientation

Career Building Workshops

Informal Graduate School Discussions

Seminar Talks

Career Building Workshops

Research Talks

Program End

Throughout: Career Building Workshops, Seminar talks, Weekend Get Togethers, and group meetings

| Activity Type | Description |
|---|---|
| Starting event | **Program Orientation**: The program held an introductory orientation session virtually in the form of a video conference that consisted of both students and mentors. |
| Professional development | **Career building workshops**: All students participated in various career building workshops; one such example was a presentation by Dr. Ondrechen on fellowship applications. **Informal Graduate school discussions**: Mentors held informal discussions on graduate school and networking for participating REU students. |
| Scientific | **Seminar talks**: PI and mentor presentations on various topics related to their research such as background on the SARS-CoV-2 targets, designing inhibitors and binders that are light activated, as well as topics related to DEI. Students additionally gave presentations on their own work or a relevant literature presentation. |
| Research | **Computationally driven structure-based ligand design:** Students took different protein targets of SARS-CoV-2 and learned how to dock libraries of compounds into these targets and to interpret the results. This was followed by short reports for researchers, with details about the findings of the best compound. In some cases, compound predictions were sent to experimentalists for synthesis and laboratory testing. Students analyzed several million different compounds against SARS-CoV-2 targets.<br><br>The different groups learned about various approaches to the analysis of protein targets, including molecular dynamics simulations and quantum mechanical modeling of SARS-CoV-2 targets and the predicted ligands. |
| Other | **Weekend Get Together**: Students would gather virtually and organize fun activities and games to promote a sense of social cohesion amongst the group. This was organized by students. |
| Ending event | **Research Talks:** Students presented a talk on their research projects and findings at the end of the summer with other REU students and faculty. |

### Research Experiences for Undergraduates in Genetic and Biochemical Mechanisms of Prokaryotic and Eukaryotic Organisms

PI: Fern Tsien & Allison Augustus-Wallace; General scientific theme: Genetics & Biochemistry

Website: https://www.medschool.lsuhsc.edu/genetics/reu.aspx

Abstract: https://nsf.gov/awardsearch/showAward?AWD_ID=1659752&HistoricalAwards

#### Programming

- The nature of research was bioinformatics, protein modeling, and data analytics.
- 25 students and 18 mentors
- Expected to work 5 days a week; no specified number of hours
- Synchronous and asynchronous components

#### Mentors

- Experienced mentors did have to adapt their projects to a virtual model; some mentors had benchwork and animal research, which they then sent their data to the students to discuss and analyze together.
- Students and mentors met at least 3 times during the week to discuss research projects.
- Diversity and Inclusion modules were required for all mentors prior to the program beginning.

#### Influence of Pandemic

- Nature of research changed in that it was genetic and biochemical wet-lab and/or computational research using in vivo and in vitro model organisms in previous years; however, it was predominately computational this iteration.
- No new mentors were recruited as a result of the pandemic.
- Students were partnered with alumni from other summer programs in the institution working with the same mentor and/or graduate students/post-docs.

#### Activities

Program Start

- CITI and KDS training modules
- Program Orientation
- Virtual Training Modules
- Professional Development Seminars
- Professional Development Seminars
- Inspiring future Scientists
- Virtual Research Poster Session
- Virtual Rewards Ceremony

Program End

Throughout: Weekly Lab Meetings, Student-Mentor Meetings, Weekly Journal Club, Weekly Departmental Seminars

| Activity Type | Description |
|---|---|
| Starting event | **Program Orientation**: This served as an introduction for students and faculty. This meeting aimed to facilitate and build the summer programs sense of community to provide more open and welcoming communication amongst students and mentors. During the program they additionally covered ground rules and expectations regarding topics such as Zoom etiquette. All students were additionally required to sign a code of conduct. |
| Professional development | **Weekly Professional Development seminars:** Each week the program hosted professional development seminars with other summer programs for approximately an hour and a half. These included how to conduct a literature search and review, scientific communication skills (presentation and manuscript writing), working as a team, time management, communicating science to non-scientists, interviewing for graduate school, professional behavior training, resume and CV writing, and career education. |
| Scientific | **Virtual Training Modules**: Prior to the program beginning all students were required to complete CITI and KDS compliance training modules which provide peer-reviewed, web-based educational courses in research practices, ethics, responsible conduct of research, and professional conduct. Additional virtual training covered topics on safety and how to maintain a lab notebook. **Software and IT support:** REU interns received IT support and had access to all necessary software to facilitate their research. |
| Research | **Mentor meetings**: Students had zoom meetings a minimum of 3 times per week with their mentors which typically lasted around two hours. During this time, mentors would discuss research projects, provide fieldwork or computational data if applicable, and answer any questions/concerns students might be having. **Weekly Lab Meetings**: These meetings typically took place once a week for about an hour; the entire lab would meet and discuss ongoing research and other relevant topics. |
| Other | **Inspiring future scientists:** Students virtually presented their projects to partnering New Orleans area middle and high school science classes to promote STEM careers while serving as role models to younger generations of potential scientists. |
| Ending event | **Virtual Poster Session**: Each student made a research poster about their respective work and submitted a 5-to-10-minute prerecorded Zoom PowerPoint presentation explaining their research, along with an abstract and headshot. **A Virtual Awards Ceremony:** This was held via Zoom, and abstracts, posters, and Zoom recordings were displayed in the REU Poster Session website. |

### Research Experience for Undergraduates in Molecular Genetics and Cell Biology

PI: Aaron Turkewitz; Scientific theme: Molecular Genetics and Cell Biology; Program Directors: Jean Greenberg, Edwin Ferguson

Website: https://mgcb.uchicago.edu/education/reu Abstract: https://nsf.gov/awardsearch/showAward?AWD_ID=1659490

#### Programming

- Nature of research was computational with mathematical modeling.
- 3 students and 2 mentors
- 40 hrs/week expected
- Synchronous and asynchronous components

#### Mentors

- One faculty member mentored two students simultaneously, but this was not as a result of the pandemic.
- Near-peer graduate students monitored progress and fielded questions.

#### Influence of Pandemic

- Nature of research differed substantially from previous years. It is normally bench-work based, with an emphasis in cellular and molecular biology approaches.
- One new mentor was recruited to the REU site this year because they had computational projects available that were suitable for undergraduates.
- An additional faculty member was recruited to give advice on computational modeling.
- Students were allowed more flexibility in scheduling their workdays than in previous years.
- Near-peers in the PhD program, some of whom were previous REUs, were liaisons between the REUs and the program leadership.

#### Activities

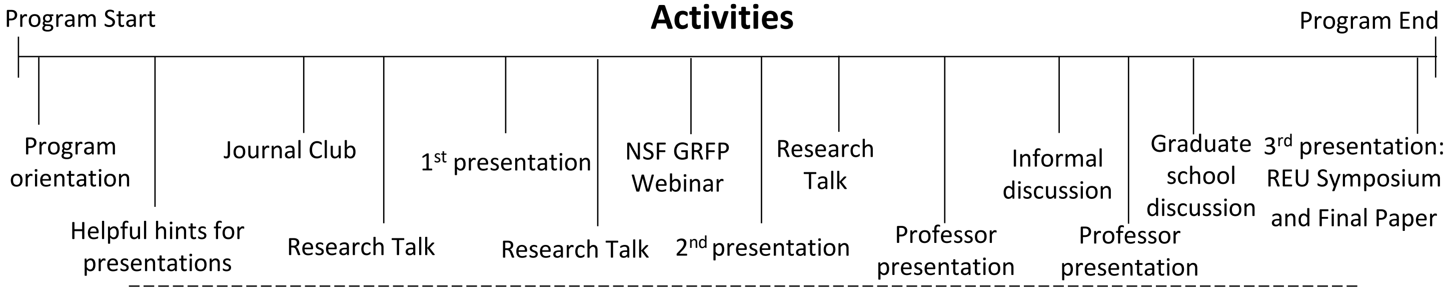


Throughout: Optional drop-ins with program leadership, Ethics workshops

| Activity Type | Description |
|---|---|
| **Starting event** | **Program orientation:** This event was held at the beginning of the program as a way introduce students to each other and program leadership. |
| **Professional development** | **Ethics workshops:** These events were held once a week throughout the first five weeks of the program. **NSF GRFP Webinar:** This event was held once during the program and included a Q&A session. **Professor presentations:** MGCB assistant professors came to speak to students twice throughout the program. **Informal discussion:** Students were given the opportunity to have an informal discussion with a scientist from the University of Chicago. **Discussion of application process for graduate school:** An MGCB professor hosted a discussion that went over the process of applying to graduate school once during the program. **Workshop on how to give a talk and how to listen to a talk:** Program leadership gave verbal discussion and provided documents to the students with guidelines. |
| **Scientific** | **Student presentations:** The first was an introduction to each project and required students to complete a one-page write-up. The second was a presentation on students' mid-program progress, and the third was the students' ending event. |
| **Research** | **Journal club:** This event was held once at the beginning of the program. **Research Talks:** These presentations were given by professors, and two former REU student (a PhD student and a recent PhD recipient), throughout the program. Each research talk was preceded by a session with program leadership to introduce the topic and discuss a paper from the speaker. |
| **Other** | **Optional drop-ins with program leadership:** Program leadership held weekly optional meetings for students to join if they wanted. **Social nights:** Near-peers organized two game nights for the REUs. |
| **Ending event** | **REU Symposium and Final Paper:** This was the final presentation students gave as a part of their participation in the program. Leadership organized a follow-up meeting several weeks after program completion to discuss the program and student plans post graduation. |

#### REU Site at The Morton Arboretum: Integrative Tree Science in the Anthropocene

PI: Chuck Cannon & Silvia Alvarez-Clare; General scientific themes: Community Ecology, Conservation & Restoration Biology, Evolution, Plant Biology, Systematics; Website: http://www.mortonarb.org/reu

Abstract: https://www.nsf.gov/awardsearch/showAward?AWD_ID=1851961&HistoricalAwards=false

##### Programming

- The nature of research centered around the following main themes: impacts of global change on tree ecology, ecosystem function, tree evolution, conservation, tree growth, management, and function in the built environment and emerging technologies in tree science.
- 8 students and 13 mentors
- 37.5 hrs/week expected
- Synchronous and asynchronous components

##### Mentors

- All students had at least two mentors.
- In most cases, an MS-level technician acted as a co-mentor for students alongside PIs at the Arboretum.

##### Influence of Pandemic

- The research themes remained the same as in previous years.
- Each mentor had to modify their project or create a new one so that students could conduct their research virtually.
- New mentors were not recruited as a result of the pandemic.
- All students worked alone (with the tech as their peer mentor) except for one group of students who worked together on separate datasets concerning the same species.

##### Activities

Program Start → Program End

- Program orientation
- Introductory writing seminar
- Science communication workshop
- Ethics workshop
- Shutdown STEM
- Applying to graduate school workshop
- Final Symposium

Throughout: Lab meetings, “Tree Talks,” Science seminars, “Meet a Scientist” events, Fun in the Field group projects

| Activity Type | Description |
|---|---|
| **Starting event** | **Program orientation:** Students were welcomed to the program, given an introductory writing seminar, and met their mentors. |
| **Professional development** | **”Meet a Scientist” events:** The program held weekly events in which guest speakers from different tree-related career paths and jobs shared their experiences with students. **Science communication workshop:** A two-hour workshop held once during the program. **Ethics workshop:** A one-hour workshop held once during the program. **Applying to graduate school workshop:** A one-hour workshop held once during the program. |
| **Scientific** | **Science seminars:** A weekly two-hour writing class with breakout room activities and assignments due each week. |
| **Research** | **Lab meetings:** Each lab had a weekly lab meeting where the REU participated. **“Tree talks”:** Weekly one-hour events that highlighted ongoing research efforts at the Morton Arboretum. |
| **Other** | **Fun in the Field projects:** Two-hour projects that consisted of tree-science projects that students completed individually outside and reported back as a group at a weekly virtual meeting. **Check-ins with program coordinator:** Weekly group meetings to cover logistics and to discuss challenges and successes. These were meant to serve as opportunities for emotional support as well. **Shutdown STEM:** A call for STEM and academic professionals to stop their usual work and spend a day learning about anti-racism and its role in science, as well as taking actions to support the Black community in STEM. |
| **Ending event** | **Final Symposium:** Symposium Keynote speakers were early-career scientists of color, including founder of Black Botanists Week who provided presentations and engagement discussions with students. Students had 12 minutes to present work and 3 minutes for questions. |

### REU Site: Global Change Ecology at the Smithsonian Environmental Research Center

PI: Alison Cawood; General scientific theme: Global Change Ecology

Website: https://serc.si.edu/internships Abstract: https://www.nsf.gov/awardsearch/showAward?AWD_ID=1659668

#### Programming

- The nature of research was mainly analysis of existing data, but there were two cases in which students were engaged in new data collection.
- 11 students and 13 mentors
- 30-40 hrs/week expected
- Synchronous and asynchronous components

#### Mentors

- Mentors did not work collaboratively to mentor the same student, but in the cases in which students worked in teams, all interns and mentors met together.

#### Influence of Pandemic

- The nature of research is typically heavily fieldwork-based, and nearly all projects include laboratory elements, but most projects this year were based on data analysis.
- No new mentors were recruited as a result of the pandemic.
- All labs were able to either adapt the original project idea to focus on existing data or developed new projects that fit within the scope of the lab, the intern's interests, and could be done remotely.
- Some students did work in teams, but not all students within a team had the same mentor.

#### Activities

Program Start

Program kick-off

Introduction to library resources

Introduction to R programming

Science communication workshops

Responsible conduct of research

Creating science videos for the public

Writing a scientific paper

Giving a scientific presentation

Exit interviews

Final presentations and video screens

Program End

Throughout: Intern social hours, Group meetings with panel discussions, Intern work hours, Office hours, Virtual SERC tours

| Activity Type | Description |
|---|---|
| Starting event | **Program kick-off:** Students were welcomed and given and introduction to SERC by the site director. Interns and mentors gave brief introductions as well. This event was 90 minutes in duration. |
| Professional development | **Introduction to R programming:** This training was a 4-part series of 3-hour workshops. **Science communication workshop:** This training was a 4-part series of 1-hour workshops. **Making a poster training. Group meetings with panel discussions:** Weekly full group meetings were facilitated by the REU site PI and included share outs from interns and a 90-minute panel discussion. Panels included SERC staff as well as outside speakers. Panel topics included Diversity and Inclusion in STEM, non-academic uses for environmental data, STEM careers, tips for grad school, and a panel of former SERC interns who had been co-authors on papers related to their REU work or who had presented their REU work at conferences. |
| Scientific | **Introduction to library resources training. Giving a scientific presentation training. Creating science videos for the public training. Writing a scientific paper training.** |
| Research | **Responsible conduct of research training:** This was a discussion-based training session. **Collaborative intern work hours:** These were held weekly. |
| Other | **Intern social hours:** These weekly events were facilitated by the intern coordinator and included games and activities. **Virtual tours of Smithsonian sites. Video assistance and presentation assistance office hours:** These started in week 8 of the program. |
| Ending event | **Final presentations and video screens:** Each intern gave a 12-minute presentation for a scientific audience and showed a 3-minute video for a public audience based on their internship work. **Exit interviews:** Students had interviews with the site PI during the last week of the internship. These were about 30 minutes in duration. |

### Rocky Mountain Biological Laboratory: Research Training in Place-Based Field Research

PI: Jennifer Reithel and Rosemary Smith; General scientific theme: Ecology and Evolutionary Biology

Website: https://www.rmbl.org/students/undergraduates-beyond/summer-education-programreu/

Abstract: https://www.nsf.gov/awardsearch/showAward?AWD_ID=1755522

#### Programming

- Nature of research was Ecology and Evolutionary Biology.
- 39 students and 16 mentors
- 40 hrs/week expected (not enforced)
- Hybrid program with most students participating in in-person activities and 6 students entirely remote

#### Mentors

- Mentors regularly collaborated, and several of the group projects depended heavily on onsite mentor and offsite mentor collaboration.
- Mentors discussed all initial project ideas with program coordinators and met with their lab teams at least once a week.

#### Influence of Pandemic

- Due to the pandemic, mentors had to adapt new projects or repurpose existing projects. They developed a hybrid on-site and remote model that allowed both students and mentors to decide how they would participate.
- Rather than have every student conduct an individual field research project as was typical in years past, some students worked on group research projects in which onsite students collected data and remote students then processed and analyzed this data.
- The lab applied for and received ROA funding which allowed for two local scientists to be recruited to serve as in-person mentors for scientists unable to be onsite for field research.

#### Activities

Program Start — Program End

- Program orientation
- Weekly workshops and panel discussions begin
- Research Proposal sessions
- Proposal writing
- Final Project Presentations

Throughout: Weekly skills workshops, panel discussions, lab group and mentor meetings, fieldwork for onsite students

| Activity Type | Description |
|---|---|
| Starting event | **Program Orientation:** The first three days of the program included an online orientation program for all 39 students. |
| Professional development | **Mentor Development:** Mentors were provided with materials from the “Entering Mentoring” program, including links to mentoring skill development. Additionally, graduate students organized a mentoring network for the program's participants. **Weekly Panel Discussions:** At least 8 panel discussions were given throughout the summer to all program participants on topics including diversity issues, Title IX, animals in research, science communication, and applying to graduate school. |
| Scientific | **Weekly Workshops:** These workshops focused on topics such as statistics, graphing, GIS, and metadata submission. |
| Research | **Proposal sessions:** Research proposals took place in sessions throughout the third week of the program for approximately 1-1.5 hours, where each student or student group would present for approximately 10-15 minutes. Approximately half the students worked in groups on research projects, where on-site group members would collect field data and remote students would then organize and analyze the data. Additionally, some students manipulated and analyzed existing datasets or samples provided by mentors. |
| Other | **Student Organized Activities:** Students organized virtual game nights and some onsite students self-organized hikes and outings (which has been typical in years past). |
| Ending event | **Final presentation:** All presentations were given virtually throughout a span of several days over a series of Zoom sessions which lasted approximately 1.5 hours each. Each student or student group was given 20 minutes to present their research projects and take questions. |

### Rosetta Commons REU: A Cyberlinked Program in Computational Biomolecular Structure & Design

PI: Jeffrey J. Gray; Program Manager: Camille Mathis; General scientific theme: Biomolecular Structure and Design

Website: https://www.rosettacommons.org/about/intern Abstract: https://nsf.gov/awardsearch/showAward?AWD_ID=1950697

https://orcid.org/0000-0001-6380-2324

#### Programming

- Nature of research was computational.
- 14 students and 14 mentors
- 37-40 hrs/week expected
- Synchronous and asynchronous components

#### Mentors

- There was one pair of mentors comprised of a graduate student/postdoc mentor and a lab PI who worked collaboratively to mentor one student.

#### Influence of Pandemic

- Nature of research did not differ from previous years; all projects are always computational.
- No new mentors were recruited as a result of the pandemic.
- Mentors did not have to develop new projects as a result of the pandemic.
- Students did not work in teams as a result of the pandemic, aside from one pair of students who worked collaboratively on their project.
- Typically, students have access to a local REU and participate in their activities, but this year they participated in virtual events hosted by the VROOM group.

#### Activities

Program Start

Program orientation

Coding Bootcamp

Student-organized social activity

JHU Scientific Seminar

Student-organized social activity

RosettaCON

Program End

Throughout: Virtual journal club, VROOM Seminars, Professional Development lessons and videos

| Activity Type | Description |
|---|---|
| Starting event | **Program orientation:** Students were welcomed and introduced to the program. It was a meet-and-greet structured event, and students participated in ice-breakers to get to know one another. It was a 2-hour long event. |
| Professional development | **Professional Development lessons and videos:** Students were given lessons on topics such as how to give an oral presentation, how to take constructive feedback, how to make a poster, how to communicate with their mentor, etc. These were held once a week and typically lasted anywhere from 30 minutes to an hour. |
| Scientific | **JHU Scientific Seminar:** A faculty member was invited to speak to students over Zoom. This event was held once and was an hour in duration. |
| Research | **Virtual Journal Club:** This is a regular part of the REU in which students present and review journals, and faculty provide a professional-development mini-lesson. The club meetings were held weekly and typically lasted an hour and a half. |
| Other | **Social activities:** Students organized these activities amongst themselves. They typically lasted about 2-3 hours and were held two times throughout the program. |
| Ending event | **RosettaCON:** This is a regular part of the REU program, but it was held virtually this year. Students attended talks and presented their posters. It occurred once at the end of the program and was held over the span of 3 days. |

### Summer Integrative Neuroscience Experience (SINE) in Jupiter

PI's: Alex Keene & Johanna Kowalko; General scientific theme: Neuroscience & Computational Biology

Website: https://www.fau.edu/jupiter/research/reu-sine/ Abstract: https://www.nsf.gov/awardsearch/showAward?AWD_ID=1852175

#### Programming

- The nature of research was computational biology and neuroscience.
- 10 students and 8 mentors
- ~40 hours/week expected
- Synchronous and asynchronous components

#### Mentors

- Mentors did not work in teams but instead worked individually with their assigned student(s). Mentors collaborated on overall goals, however, not on research specifics.
- PIs checked in with mentors every two weeks.

#### Influence of Pandemic

- The intended nature of research was immersive wet lab work with a focus on molecular and behavioral neuroscience, however, due to COVID projects had to be done virtually and all research became computational.
- As a result of COVID restrictions, the mentor pool shrank leaving some mentors with two students.
- This was the first year of research, however, approximately half the intended projects were altered as a result of the pandemic, and half the projects ran analysis on ongoing projects.
- Each student had their own project but would often work in teams to discuss and analyze these projects in hopes of creating social bonds.

#### Activities

Program Start — Program End

Mentor & Group Orientations → Professional Development workshops → Professional Development workshops → Final Research Talks

Throughout: Structured Coursework, guest speakers, career development panels, research seminars, PI check-ins with mentors , student data-analysis, scientific communication, journal club, and microscopy coursework.

| Activity Type | Description |
|---|---|
| Starting event | **Mentor Orientation**: Prior to the programs start, there was a one-hour discussion with mentors where they covered objectives and potential challenges for the upcoming summer program. In addition, the PIs met with incoming students in 2 one-hour meetings to discuss the program. On the first day of orientation, all mentors (faculty and staff) participated in conversations about goals for the summer. The PIs moderated a conversation about engaging in scientific research and the career path of faculty mentors. |
| Professional development | **Career development panels**: The program organized a multitude of professional development events such as bringing in graduate students, groups of faculty that were part of graduate admissions, and other guests to speak and give advice in the place of coursework on some mornings. The program hosted 6 **professional development workshops:** Most of them involved outside speakers with expertise in the field. Each was two hours long and the topics covered include an introduction to graduate school, the application process, how to evaluate potential graduate school programs and mentors, how to talk to a potential mentor, graduate school interviews, as well as what happens after graduate school. |
| Scientific | **Research seminars:** The four days of the week that the group formally met each afternoon consisted of an hour and a half allocated towards attending presentations from neuroscientists or having a paper discussion over upcoming speakers, research papers, and scientific work. **Scientific Communication:** Students participated in journal club and discussed scientific presentations, ethics, and the peer review process for approximately four hours per week. **Journal Club:** On two days of the week the group went over a paper by the following day's guest speaker; students devoted eight hours per week to journal club. Then students discussed the paper with the scientist who was lead or corresponding author. Students were guided to develop questions they could ask the visiting speakers. In total, there were 16 guest speakers that ranged from graduate students to faculty. **Microscopy course:** This two-hour per week course went over microscopy basics, confocal imaging and data analysis. The students participated in virtual imaging exercises, where samples were imaged in real time. |
| Research | Students were encouraged to spend approximately four hours each day working on data analysis for their host laboratories. The PIs checked on student progress on a biweekly basis. In addition, the **scientific communications** course was used for group discussion of lab projects. All students participated in **weekly lab meetings** and presented at least once during the semester. |
| Other | PI's encouraged students to discuss diversity within science and had regular conversations about racial injustices and inequality. |
| Ending event | **Research Talks**: At the end of the program all students presented a 10 to 15-minute talk on their research. The event was attended by mentors and members of the community affiliated with the program. |

#### Teaching plant and agricultural phenomics through unPAK

PDs: Matt Rutter, Allan Strand & Courtney Murren, Evaluator Danielle Jensen-Ryan; General scientific themes: Plant science, Genotype to Phenotype, Phenomics, Agriculture, Evolutionary Ecology; Website: arabidopsisunpak.org

##### Programming

- The nature of research was computational and experiment-based.
- 11 students and 4 mentors and additional home institution mentors
- 37.5 hrs/week expected
- Synchronous and asynchronous components

##### Mentors

- Mentors did work collaboratively with other mentors to guide and advise students, but this was not as a result of the remote circumstances.

##### Influence of Pandemic

- Nature of research did not change significantly, but it was more computational than in previous years.
- The senior scientist mentorship team was the same as in 2019. However, the Zoom and Slack-based communication enabled home-institution mentors from which summer participants were recruited to be connected during the program.
- Mentors developed an entirely new plant-growth kit for a phenotyping project, along with training videos.
- Students did work in teams or pairs, but not as a result of the pandemic.

##### Activities

Program Start

Opening discussion

R workshop

Plant science database workshop

Ethics workshop

The process of writing

R workshop

Workshop on library research

R workshop

E-fieldtrip to Selma and the Jackson Foundation Museum

Final presentations

Program End

Throughout: Guest seminars, FunPAK events, Invited e-seminars

| Activity Type | Description |
|---|---|
| **Starting event** | **Opening discussion of expectations and discussions about organizing one's remote workday:** These discussions were held briefly and regularly as well as at the beginning of the program. |
| **Professional development** | **R workshops:** These two-hour workshops were held three times throughout the program and went over topics such as data management, graphing, and statistical approaches. **Ethics workshop:** This one-hour event was held once during the program. **Guest seminars:** Speakers from diverse career stages, institutions, diverse racial and ethnic backgrounds, and from both the US and other countries, presented to students. There were Q&As on plant science topics, and speakers also spoke of their journey as a scientist. CV development and discussion of the broad framework of USDA program and agricultural careers. |
| **Scientific** | **The process of writing:** This workshop was held formally once during the program and was an hour in duration and was revisited periodically throughout the program. |
| **Research** | **Workshop on plant science databases:** This one-hour event was held once during the program. **Workshop on library research:** This one-hour workshop was held once during the program. |
| **Other** | **FunPAK:** These one-hour events were held weekly and consisted of games, art, and sharing recipes. **Invited e-seminars:** These one-hour seminars were held once a week. **e-fieldtrip to Selma and the Jackson Foundation Museum:** Students virtually visited the house where civil rights and voting rights leaders (including Dr. Martin Luther King) planned and coordinated the Selma to Montgomery March (jacksonfoundationandmuseum.com). The conversation focused on the civil rights era, social justice, antiracism, the impact of young people on the movement, and the connection to voting rights of today. This fieldtrip contributed to students expressing their views and discussing civil rights and social issues. |
| **Ending event** | **Final presentations:** These presentations were day-long oral presentations via Zoom. Team presented outcomes of the two cohort/team projects (~15 minutes followed by 5min Q&A) and individual/pair projects (~10 minutes followed by ~2 min Q&A). Invited guests included home mentors and peers so students could get feedback on presentations. |

### Training and Experimentation in Computational Biology (TECBio) at the University of Pittsburgh

PI: Joseph Ayoob; General scientific theme: Computational and Systems Biology

Website: www.tecbioreu.pitt.edu   Abstract: https://www.nsf.gov/awardsearch/showAward?AWD_ID=1659611

#### Programming

- The nature of research was computational structural biology, cellular and systems biology, genomics and bioinformatics, computational drug discovery, and bioimage informatics.
- 16 students and 15 mentors
- 40 hrs/week expected
- Synchronous and asynchronous components

#### Mentors

- There was one experienced mentor, whose background is mainly experimental. They collaborated with another TECBio mentor to build a model with their student instead of gathering new experimental data.

#### Influence of Pandemic

- Nature of research did not differ substantially from previous years.
- New mentors were recruited to the site, but they were not recruited as a result of the pandemic.
- The workload for students was slightly decreased; some individual activities became group projects.
- Some students worked in teams, which was new this year; two mentors also worked collaboratively to mentor the same student.
- Program leadership incorporated the use of virtual reality technology to help foster interpersonal interactions among students and program directors.

#### Activities

Program Start → Program End

- Program orientation
- Entering research training for mentees
- Research roundtable
- Ethics forum
- Proposal writing
- Research roundtable
- Virtual reality poster session and Zoom symposium

Throughout: Student leadership teams, research and career seminars, informal meetings with PI, journal club, TGIF

| Activity Type | Description |
|---|---|
| Starting event | **Program orientation:** Students were welcomed and introduced to the program. As this was the first meeting together, the program made introductions by forming small groups of students to get to know each other in Zoom breakout rooms, and program and student expectations were discussed in the whole group. |
| Professional development | **Entering research training for mentees:** The PI developed a 5-session mini-arc of this curriculum. In these sessions, students discussed their excitement and concerns about their research experience, ways to align their goals and expectations with those of their mentors, challenges facing diverse teams, and ways to handle difficult situations and tasks by relying on the program infrastructure and mentoring circle as well as cultivating their own self-efficacy. **Ethics forum**: In small groups, students investigated an instance of scientific misconduct, prepared a discussion on the topic, and presented it to students in other summer programs. Graduate students served as mentors for the groups and help them prepare their discussions. |
| Scientific | **Research and career seminars:** The program hosted a weekly one-hour seminar series for our students. These sessions introduced students to the various scientists in the field, their research, and their career arcs. **Journal club:** This event was held weekly; students participated in teams of 2-3. |
| Research | **Proposal writing:** Students submitted a 4-page proposal outlining their research goals/hypotheses, methods, alternatives, and significance of expected results by the end of the fifth week. Students wrote these independently with input and guidance of their mentors. **Research Roundtables:** Twice during the summer, the program hosted discussions where students briefly described their research project/question, its importance, the approach, and any findings they made and/or challenges they are facing. |
| Other | **Student leadership teams:** Students each served on two of the following teams to address the specific goals in consultation with a graduate student advisor and the PI: mentoring team, t-shirt team, ambassadorship team, social team. **TGIF (TECBIO Games, Interviews, and Fun):** These events were networking/social gatherings that ran throughout the program at the end of the week. |
| Ending event | **Virtual reality poster session and Zoom symposium:** Students presented their work orally on the second to last day of the program in Zoom, which accommodated the attendance of their friends and family. On the last day, the program hosted a poster session in VR. |

### A Transformative Approach for Engaging First-Generation Underrepresented Minorities in a Research Experience

PI: Bob Kao; General scientific theme: Microbial & Cell Biology of Animal Development

Website: https://www.heritage.edu/academic-paths/special-programs/hu-nsf-reu/

Abstract: https://www.nsf.gov/awardsearch/showAward?AWD_ID=1852032

#### Programming

- The nature of research was applied bioinformatics and computational biology.
- 11 students and 6 mentors
- 40 hrs/week expected
- Synchronous and asynchronous components

#### Mentors

- Mentors and the PI collectively helped plan new and adapted research experiences.
- Mentors collectively worked together using a sociocratic model to design new projects for virtual research experiences and collaborated on projects frequently during community of scholars discussions.

#### Influence of Pandemic

- The nature of research in the past typically included in-person lab studies combined with field studies, however, no in-person lab work nor field studies were permitted due to stay-at-home orders.
- The lab collectively decided to use a systems thinking community framework to plan for an applied bioinformatics remote research summer experience.
- No new mentors were recruited as a result of the pandemic.
- The number of undergraduate participants was increased so that undergraduates could work in teams in the new virtual format.

#### Activities

Program Start

Virtual pre-REU workshop

Research Overview

Intro to Image J Data Analysis

Discussions on Black Lives Matter and topics in on inclusion, equity, diversity

Ethics of Image Analysis

Refining research proposals and review articles

Community of Scholars Presentations using Padlet

Program End

Throughout: Community of scholars discussions, guest speakers, professional development networking, and webinars

| Activity Type | Description |
|---|---|
| Starting event | **Virtual pre-REU Workshop:** The PI and mentors had a virtual workshop for all REU participants for 2 hours each day over the course of three days. During the workshop, adjustments were collectively made to in-person lab techniques, new research proposals were adapted, and participants worked in groups to present a mini research proposal. |
| Professional development | **SACNAS webinars:** Throughout the ten-week virtual research program, undergraduates attended webinars on Writing an Individual Development Plan and Harmony in Work and Life. Additionally, three guest speakers were brought in to discuss their own experiences in the field of science and provide input on topics such as communication in science, graduate school, and career paths. |
| Scientific | **Community of scholars discussions:** These discussions occurred 4-5 days a week and were typically 2 hours each. Researchers would cover how the applied bioinformatic technique fits into the fields of cell and microbial developmental biology and how to define research questions. This was followed by a step-by-step walk through of ImageJ and how to interpret data from research papers. Additionally, this was typically followed up with an interactive Q&A. The discussion would then end by summarizing the approach and linking it back to the big picture of the field. |
| Research | Participants were given the option to work in teams to create applied bioinformatics posters which were then displayed in a virtual gallery. |
| Other | Student pop-up hours via Zoom were provided throughout the 10-week summer research experience to go over undergraduates' topics for their respective research proposals and posters. |
| Ending event | The program collectively shared research proposals, topics for review articles, and posters on applied bioinformatics using Padlet and Zoom discussions. |

#### FOCUS GROUP QUESTIONS

**Student midpoint**

1. Briefly tell me about your experience this summer so far – who are you working with? What are you working on?
2. What is one thing about the program that is going well for you?
3. What is one suggestion you have for improving the program or your research experience?
4. Anything else you think it would be important for me or other people involved in the program to know?

**Student endpoint**

1. What is the most important thing you got out of the experience?
2. As a reminder, here are some of the things you highlighted as going well the last time we spoke: 30,000-foot view of what was going well blurb. Anything to add to that?
3. As a reminder, here are some of the things you highlighted as areas of improvement: 30,000-foot view of areas of improvement blurb. Anything to add to that? Are these still areas of improvement?
4. Is there any aspect or part of the program that you would recommend dropping?
5. If you could give one piece of advice to a student participating in a remote REU in the future, what would it be?
6. If you could give one piece of advice to a mentor in a remote REU in the future, what would it be?
7. Anything else that would be important for us to know?

**Mentor midpoint**

1. What are one or two things that are going well in the program and/or with your undergraduate researcher thus far?
2. What are one or two suggestions for improving the program and/or your work with your undergraduate researcher for the remainder of the summer?
3. Anything else you think it would be important for me or other people involved in the program to know?

#### SURVEY MEASURES

##### Synchronous vs. asynchronous programming

**Item:** Were events in your program (workshops, guest speakers, etc.) mostly synchronous (live) or asynchronous (recorded)? **Response options:** (1) Entirely synchronous, (2) Mostly synchronous, (3) Both synchronous and asynchronous, (4) Mostly asynchronous, (5) Entirely asynchronous, (6) I prefer not to respond.

**Relationship quality** (Ragins & Cotton, 1999)
**Items:** Think of the person that mentored you most in your research this summer. With this person in mind, please indicate the extent to which you agree with the following statements… **Response options:** (1) Strongly disagree, (2) Moderately disagree, (3) Slightly disagree, (4) Slightly agree, (5) Moderately agree, (6) Strongly agree, (7) I prefer not to respond.

1. My mentor is someone I am satisfied with.
2. My mentor has been effective in their role.
3. My mentor always met my needs.
4. My mentor never disappointed me.

**Connectedness (**Rovai, 2002)
With your summer research program in mind, please indicate the extent to which you agree or disagree with the following statements… **Response options:** (1) Strongly disagree, (2) Moderately disagree, (3) Slightly disagree, (4) Slightly agree, (5) Moderately agree, (6) Strongly agree, (7) I prefer not to respond. Note: R indicates an item that should be reverse scored.

1. I feel that students in this program care about each other
2. I feel connected to others in this program
3. I do not feel a spirit of community (R)
4. I feel that this program is like a family
5. I feel isolated in this program (R)
6. I trust others in this program
7. I feel that I can rely on others in this program
8. I feel that members of this program depend on me
9. I feel uncertain about others in this program (R)
10. I feel confident that others will support me
